# Cell-type-specific plasticity shapes neocortical dynamics for motor learning

**DOI:** 10.1101/2023.08.09.552699

**Authors:** Shouvik Majumder, Koichi Hirokawa, Zidan Yang, Ronald Paletzki, Charles R. Gerfen, Lorenzo Fontolan, Sandro Romani, Anant Jain, Ryohei Yasuda, Hidehiko K. Inagaki

**Affiliations:** Max Planck Florida Institute for Neuroscience, Jupiter, FL 33458, USA; National Institute of Mental Health, Bethesda, MD 20814, USA; Turing Centre for Living Systems, Aix- Marseille University, INSERM, INMED U1249, Marseille, France; Janelia Research Campus, HHMI, Ashburn VA 20147, USA

## Abstract

Neocortical spiking dynamics control aspects of behavior, yet how these dynamics emerge during motor learning remains elusive. Activity-dependent synaptic plasticity is likely a key mechanism, as it reconfigures network architectures that govern neural dynamics. Here, we examined how the mouse premotor cortex acquires its well-characterized neural dynamics that control movement timing, specifically lick timing. To probe the role of synaptic plasticity, we have genetically manipulated proteins essential for major forms of synaptic plasticity, Ca^2+^/calmodulin-dependent protein kinase II (CaMKII) and Cofilin, in a region and cell-type-specific manner. Transient inactivation of CaMKII in the premotor cortex blocked learning of new lick timing without affecting the execution of learned action or ongoing spiking activity. Furthermore, among the major glutamatergic neurons in the premotor cortex, CaMKII and Cofilin activity in pyramidal tract (PT) neurons, but not intratelencephalic (IT) neurons, is necessary for learning. High-density electrophysiology in the premotor cortex uncovered that neural dynamics anticipating licks are progressively shaped during learning, which explains the change in lick timing. Such reconfiguration in behaviorally relevant dynamics is impeded by CaMKII manipulation in PT neurons. Altogether, the activity of plasticity-related proteins in PT neurons plays a central role in sculpting neocortical dynamics to learn new behavior.

## Introduction

Neural computations are mediated by time-varying and coordinated spiking activity across a population of neurons, referred to as neural dynamics. For example, during the planning of volitional movement, the premotor cortex exhibits slowly varying neural activity that determines the type and timing of upcoming movement, referred to as preparatory activity^1,2^. Previous research has shown flexible reconfiguration of dynamics, including preparatory activity, during motor learning^3–16^. But the neural mechanisms that reshape the neocortical dynamics to enable acquisitions of new behavior remain elusive.

Network architectures, i.e., synaptic connections, constrain neural dynamics^17^. Therefore, altering specific connections in the network architecture through synaptic plasticity is the primary theory for learning^12,14,18^. First, experience- and learning-dependent synaptic plasticity has been observed across many brain areas^7,19–27.^ Second, essentially all neurons are equipped with molecular pathways that mediate plasticity^28–32^. Third, manipulations of synaptic plasticity influence learning, although this has primarily been investigated in the hippocampus, amygdala, and cerebellum^33–38^ and far less in the neocortex^11,39–41^.

The neocortex contains diverse neuronal cell types across layers, characterized by unique gene expression profiles and anatomical features^42–44^. These cell types often carry distinct information and contribute to different aspects of neural dynamics and behavior in expert animals performing tasks^45–48^. The function of synaptic plasticity during learning may also vary across neocortical cell types to shape these cell-type-dependent dynamics and behavior.

To probe the role of synaptic plasticity across cell types during motor learning, we performed a series of acute genetic manipulations of proteins required for synaptic plasticity. We studied learning of motor timing in mice, focusing on the premotor cortex, as it provides several advantages. First, neocortical dynamics in the premotor cortex and their causal roles on behaviors, including timing behavior, have been well-established in expert animals^1,2,49–53.^ In addition, the premotor cortex has been implicated in learning across tasks and species^10,54–56^, making it an ideal site to examine the function of plasticity in reconfiguring dynamics during learning. Finally, animals are adept at learning motor timing, making it a quick and robust system to study learning. Leveraging this model system with molecular manipulations and high-density electrophysiology, we identified the key cell type required to shape behaviorally relevant neocortical dynamics and timing of action.

## Results

### CaMKII activity in ALM is necessary to learn new lick timing

We developed an operant motor timing task, in which water-restricted mice learn to delay their lick time in order to acquire water rewards (Fig.1a and Methods). Trial onset was signaled by an auditory cue (3 kHz tone, 0.6 s), and a lick after a following unsignaled delay epoch was rewarded (rewarded trials). An early lick during the delay epoch aborted the trial without a reward. Training proceeded in two stages. First, mice were trained to lick after the cue onset with a minimal delay (0.1 s, ‘cue association’). Second, the delay duration was gradually increased (‘delay training’; reaching criterion performance, 30% rewarded trials in the last 100 trials with a given delay duration, resulted in a delay increase of 0.1 s; Methods). Following this protocol, mice learned to delay lick timing within and across sessions (Fig. 1b; lick time reached 1.20 ± 0.31 s, mean ± SEM, in 6 days of delay training, n = 6 mice). High-speed videography (300 Hz) revealed that animals moved their jaw and tongue before the full tongue protrusion (Extended Data Fig. 1a-f). Yet, the delayed lick time after training was primarily due to the withholding of orofacial movement (Extended Data Fig. 1a-f).

**Figure 1.**
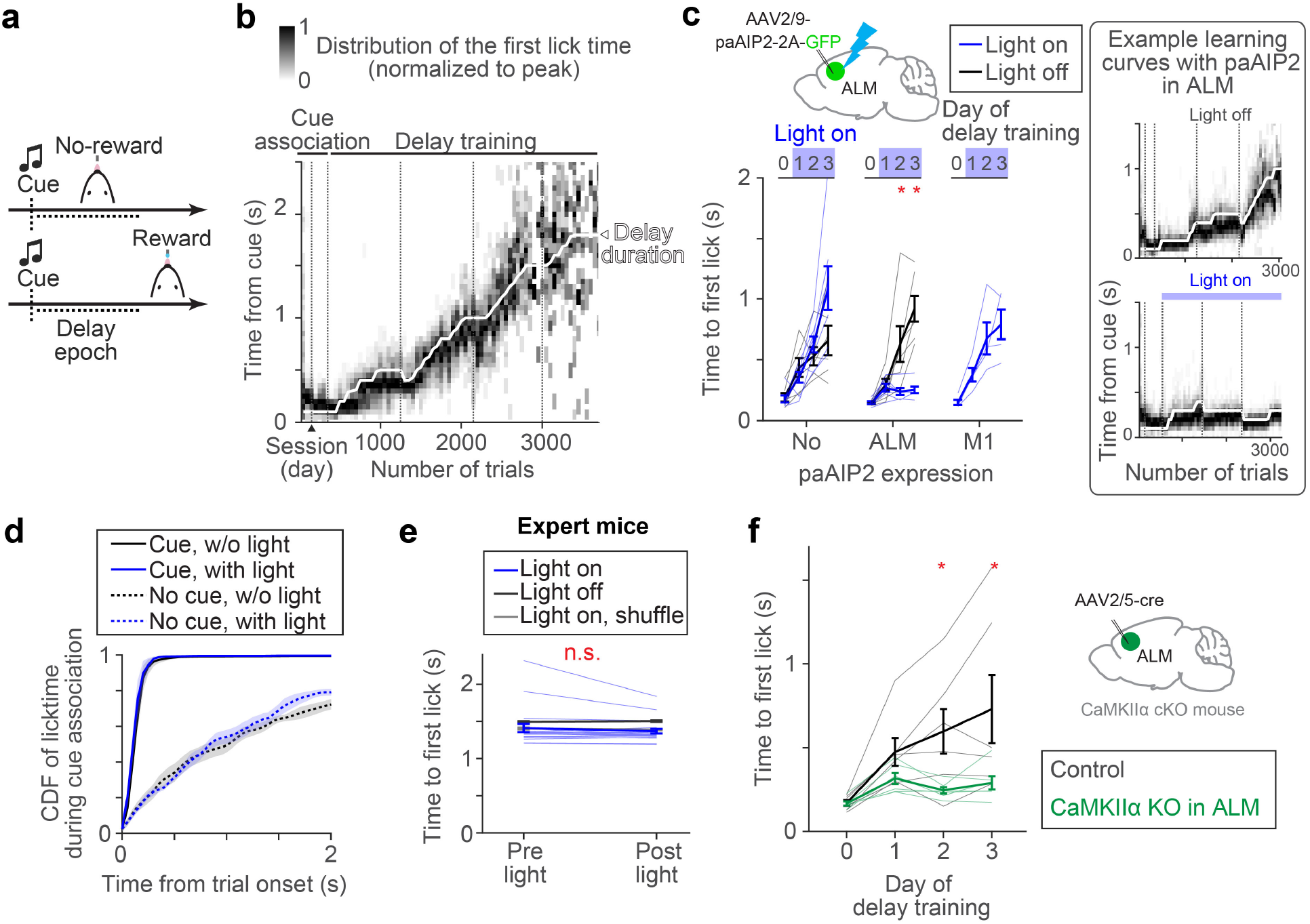
CaMKII activity in ALM is required for delay learning. **a.**Task design. **b.** Example learning curve. Distributions of lick time per trial bin are shown (50 trials; Methods). Vertical dotted lines separate sessions. **c.** paAIP2 manipulation during learning. The blue light was on during the delay training but not cue association. Day 0, the last day of cue association. The median lick times of the last 100 trials are shown. Thick lines, mean ± SEM. Thin lines, individual mice (n > 5 for ‘ALM’ and ‘No’ per condition, and n = 4 for ‘M1’). *: *p* < 0.05 (bootstrap followed by *Bonferroni* correction). See Extended Data Table 3 for the exact n and *p*-values. Right, example learning curves of mice with paAIP2 expression in ALM (top, light off; bottom, light on). The format is the same as in **b**. **d.** Distribution of first lick time on the second day of cue association. ‘Cue’ and ‘No cue’, trials with and without a cue, respectively (Methods; cue starts at the trial onset in cue trials). Different distributions between cue and no cue trials indicate successful cue association. Shade, SEM. *pcue with vs. w/o light* = 0.571 (ranksum test, n = 3 and 5 mice, light on and off respectively). **e.** ALM paAIP2 manipulation in expert animals. Comparing the first lick time within a session (100 trials before and after the onset of blue light). Thick lines, mean ± SEM. Thin lines, individual sessions (n = 4 mice, 20 sessions). *p* = 0.117 (bootstrap). See Methods for the shuffle procedure. **f.** Knocking out CaMKII⍺ expression in ALM. Thick lines, mean ± SEM. Thin lines, individual mice (n = 6 mice per condition). *: *p* < 0.05 (bootstrap followed by *Bonferroni* correction).

Next, we examined whether synaptic plasticity in the anterior-lateral motor cortex (ALM; AP 2.5mm ML 1.5 mm from Bregma), a premotor cortical area responsible for orofacial movement^5,57^, is required for this learning. To this end, we blocked the activity of CaMKII, a Ca^2+^-dependent kinase required to induce major types of synaptic plasticity and learning^27,29,30,32,33^. A transient manipulation of CaMKII activity using a genetically-encoded photoactivatable competitive inhibitor of CaMKII, paAIP2, blocks the induction of synaptic plasticity^37^ in the presence of blue light without influencing the excitability of neurons (Extended Data Fig. 2a-f).

Control mice without paAIP2 expression learned new lick timing regardless of blue light illumination of ALM (470nm, 0.5mW, 0.2 Hz; Fig. 1c, first column; Methods). In contrast, AAV-mediated bilateral paAIP2 expression in ALM blocked learning when blue light was illuminated during the delay training, but not in the absence of blue light (Fig. 1c, second column; Extended Data Table 3; Extended Data Table 1 and 2). Notably, during light illumination, mice continued licking after the cue without a change in the variability of lick time (Fig.1c and Extended Data Fig 1g-j). This implies that paAIP2 manipulation does not block animals’ ability to explore different lick times across trials (which is a prerequisite for reinforcement learning^58^). Instead, the manipulation interferes with the ability to directionally shift the distribution of lick times, presumably guided by rewards. Similar manipulation in M1 (AP 0.0mm ML 1.5 mm from Bregma) did not block learning (Fig. 1c, third column), implying that CaMKII activity in ALM is required for delay learning.

The ALM paAIP2 manipulation did not interfere with the cue association (Fig. 1d). In expert mice trained for two weeks with a fixed delay at 1.5 s (Methods), paAIP2 manipulation did not alter the distribution of lick timing, implying that CaMKII activity in ALM is not required for mice to execute learned delayed licks (Fig. 1e). Electrophysiological recording of ALM in expert mice confirmed that paAIP2 manipulation does not directly perturb ongoing spiking dynamics during behavior (Extended Data Fig. 2g-o). Altogether, CaMKII activity in ALM is specifically required for learning new lick timing but not for the initial cue association, the execution of learned action, or ongoing spiking activity.

CaMKII⍺ isoform is required for synaptic plasticity and learning, while other CaMKII isoforms are implicated in different cellular functions^30,59^. Since paAIP2 binds to the kinase domain homologous across the CaMKII family^37^, paAIP2 likely blocks all of them. To test whether CaMKII⍺ in ALM is required for learning, we acutely knocked it out in adult ALM by injecting AAV-hsyn-cre in CaMKII⍺ conditional knockout mice^60^ (Fig. 1f and Extended Data Fig. 3c). These mice learned cue association, yet could not learn to delay lick timing (Fig. 1f), consistent with the paAIP2 manipulation. Altogether, learning new lick timing requires CaMKII⍺ activity in ALM.

### Synaptic plasticity proteins in PT but not IT cells are required for motor learning

The major excitatory cell types in the premotor cortex include IT and PT neurons, projecting within and outside the telencephalon, respectively (Fig. 2a). Both of these cell types highly express CaMKII⍺ in ALM^43^ (Extended Data Fig. 3). The key question is whether synaptic plasticity in these diverse populations has redundant or specialized roles in learning. We generated an AAV expressing paAIP2 in a cre-dependent manner to manipulate CaMKII activity in distinct neocortical cell types (Fig. 2b). Strikingly, paAIP2 manipulation of PT neurons in ALM completely blocked delay learning without affecting the distribution of cue-triggered lick, similar to the bulk manipulation across cell types (Fig. 2b and c; Extended Data Fig.1). We observed consistent behavioral effects whether we labeled PT neurons using Sim1-cre KJ18 transgenic line^61^ or manipulated individual PT subtypes using retrograde AAV (PT_upper_ and PT_lower_ neurons with distinct subcortical projection^47^; 2700 ± 208 manipulated cells/animal, mean ± SEM; Fig.2c and Extended Data Fig. 4). In contrast, paAIP2 manipulations in IT neurons (layer 2/3 and 5 IT neurons) did not block learning (Fig. 2c).

**Figure 2.**
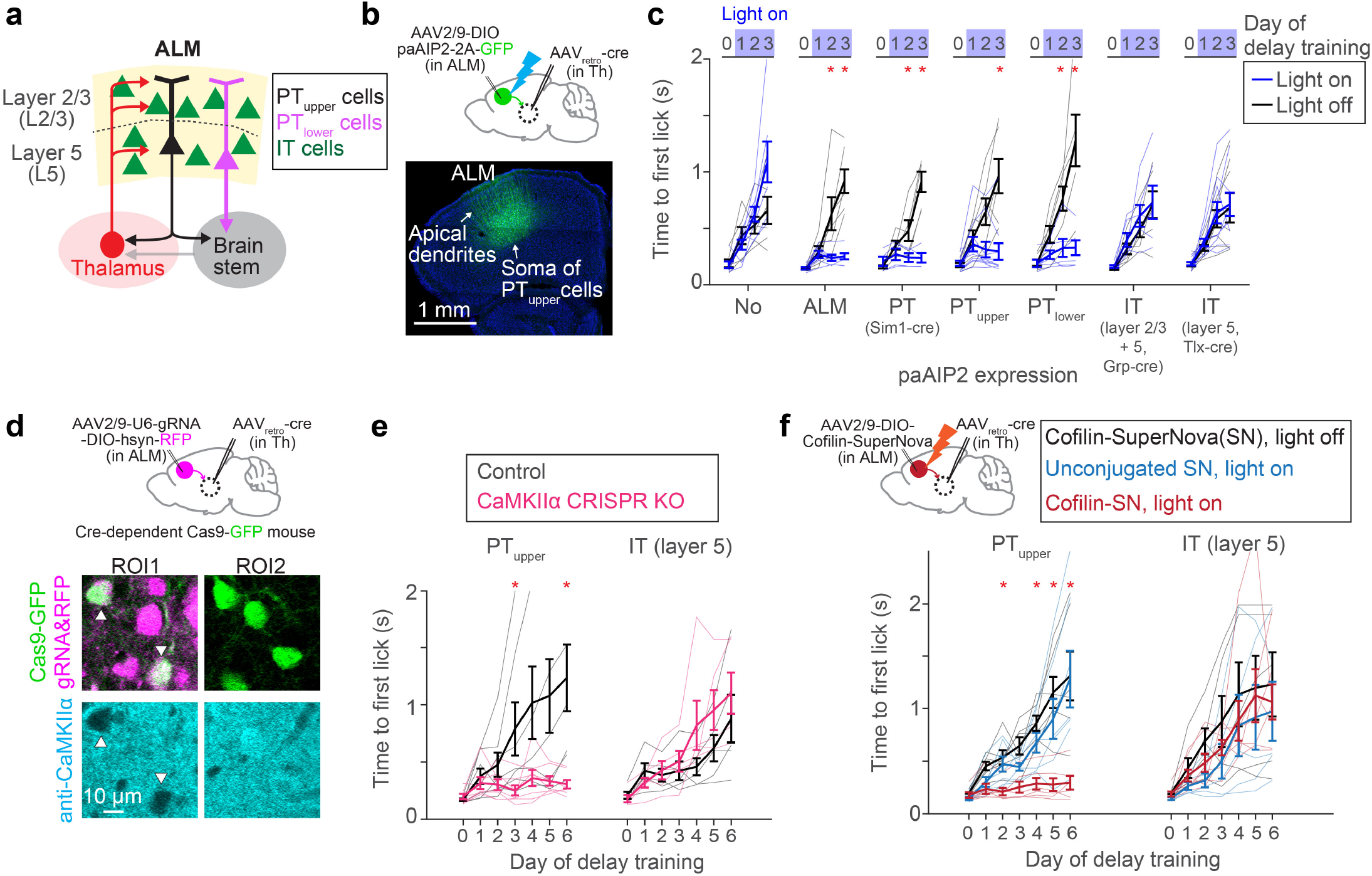
CaMKII activity in PT neurons is required for delay learning. **a.** Network architecture and cell types in ALM. **b.** Expression of paAIP2 in ALM PTupper neurons. **c.** The effect of paAIP2 manipulation in distinct cell types in ALM. ‘No’ and ‘ALM’ are duplicated from Figure 1c for comparison purposes. Thick lines, mean ± SEM. Thin lines, individual mice (n > 3 mice per condition). *: *p* < 0.05 (bootstrap followed by *Bonferroni* correction; Extended Data Table 3). **d.** Top, cell-type-specific KO of CaMKII⍺ using CRISPR/Cas9. Bottom, immunohistochemical validation of loss of CaMKII⍺ protein expression in PTupper neurons. **e.** The effect of CaMKII⍺ KO in PTupper and IT neurons. Thick lines, mean ± SEM. Thin lines, individual mice (n > 4 mice per condition). *: *p* < 0.05 (bootstrap followed by *Bonferroni* correction; Extended Data Table 3) **f.** The effect of inactivating Cofilin in PTupper and IT neurons. Thick lines, mean ± SEM. Thin lines, individual mice (n > 4 per condition). *: *p* < 0.05 for both Cofilin-SN light on vs. off comparison and Cofilin-SN light on vs. unconjugated light on conditions (bootstrap followed by *Bonferroni* correction; Extended Data Table 3).

To test the necessity of the CaMKII⍺ isoform across cell types, we acutely knocked out CaMKII⍺ using CRISPR/Cas9. We injected AAV expressing guide RNA against CaMKII⍺^30^ in ALM of adult mice expressing Cas9^62^ in PT_upper_ or IT neurons (Methods). After a month of AAV injection, a significant proportion of cells that co-express Cas9 and guide RNA lost CaMKII⍺ protein expression (Fig. 2d and Extended Data Fig. 3). PT_upper_-specific knockout of CaMKII⍺ blocked delay learning without any effect on cue association, whereas IT-specific knockout did not affect learning, consistent with the paAIP2 manipulations (Fig. 2e).

As an independent approach to manipulate synaptic plasticity, we inactivated Cofilin, a protein required for actin remodeling during major forms of synaptic plasticity^38^. To this end, we leveraged Cofilin conjugated with SuperNova (Cofilin-SuperNova), a monomer photosensitizing fluorescent protein for Chromophore Assisted Light Inactivation (CALI)^38^. PT_upper_-specific expression of Cofilin-SuperNova and orange light illumination (595 nm, 0.75 mW, 1 min every 10 min) during the delay training blocked learning without affecting cue-triggered licks (Fig. 2f). In contrast, no illumination or unconjugated SuperNova with illumination did not affect learning, implying that CALI-mediated inactivation of Cofilin blocked the learning. Consistent with the CaMKII manipulation, manipulating Cofilin in IT neurons did not affect learning. Altogether, three cell-type specific acute genetic manipulations imply that synaptic plasticity in PT but not IT neurons is necessary to learn new lick timing.

### Evolution of ALM preparatory dynamics during delay learning

Since preparatory activity determines the upcoming actions^2^, ALM preparatory dynamics are likely tailored during learning new lick timing. To characterize preparatory activity in our behavioral task, we performed acute high-density extracellular electrophysiological recordings in ALM during learning and in expert mice (Fig. 3 and Extended Data Fig. 5; with rigorous quality control in spike sorting, Extended Data Fig. 6). Among the 3203 putative pyramidal neurons we recorded from 30 mice, we focused on 1613 neurons with preparatory activity (defined as neurons with significant activity between cue and lick, signed-rank test, *p* < 0.05; Methods; Extended Data Fig. 7 and Table. 3).

**Figure 3.**
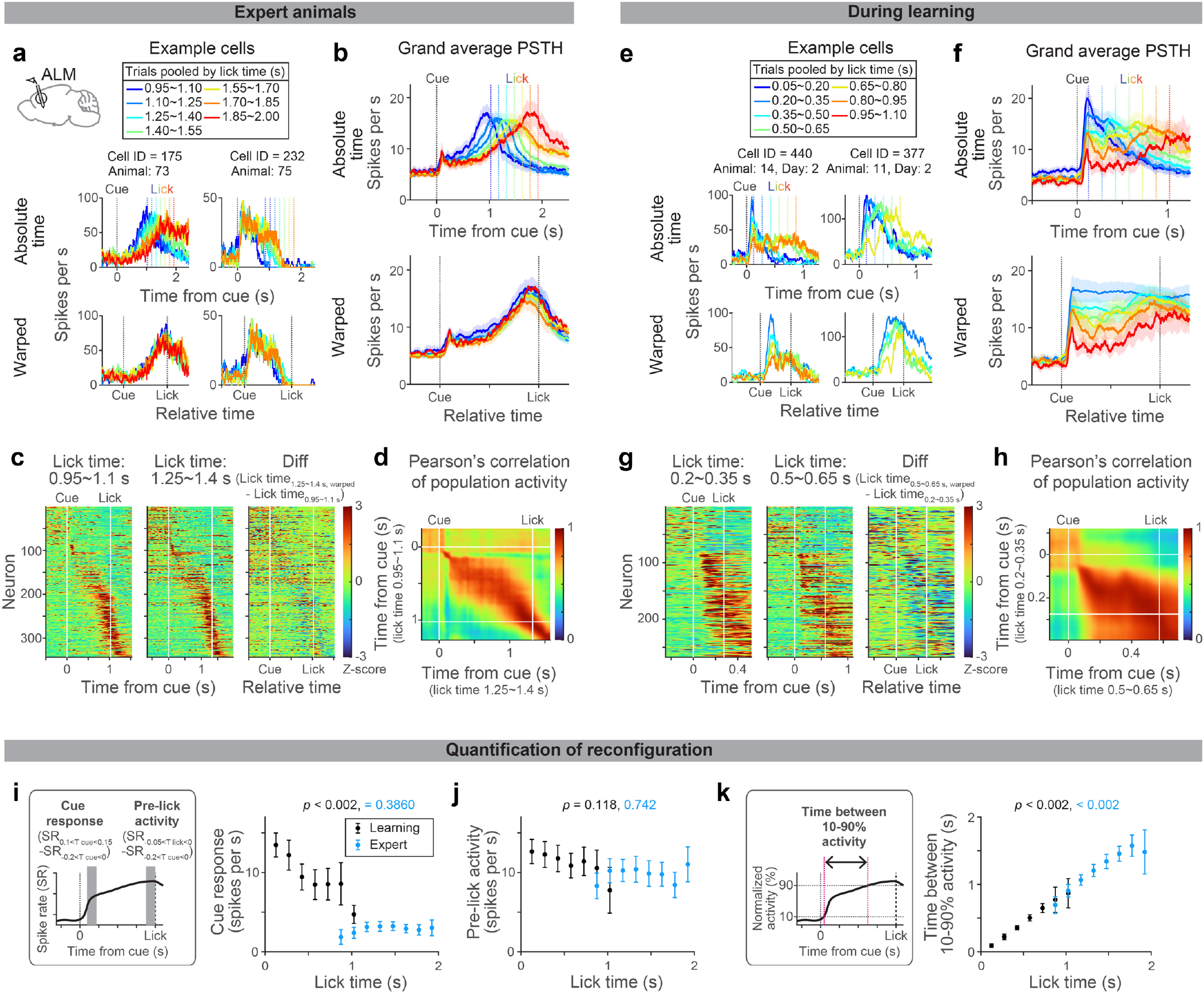
Evolution of ALM dynamics during delay learning. **a.** Spiking activity of example neurons in expert mice. Lines with different colors indicate the mean spike rate of trials with different first lick times (vertical dotted lines in the same color indicate the corresponding lick time). Top, activity in absolute time. Bottom, activity in relative time, following temporal warping (all trials were warped to have identical lick time; Methods). **b.** Grand average peri-stimulus time histogram (PSTH) of positively-modulated ALM preparatory neurons (cells with the mean spike rate between cue and lick significantly higher than the baseline). The same format as in **a**. Shade, SEM. Trial types with more than 50 neurons are shown. See Extended Data Table 3 for the number of neurons analyzed. **c.** Z-scored spiking activity of preparatory ALM neurons in trials with first lick time within 0.95-1.1 s (first column) and 1.25-1.4 s (second column). All preparatory neurons with these two trial types are shown (n = 344 cells). The difference between the first and second columns after temporal warping (third column). **d.** Pearson’s correlation of population activity between two trial types shown in **c**. **e-h.** Same as in **a-d** for during learning. **g-h**: N = 269 cells. Note that many cells show a decrease in spiking activity as mice lick later (third column), unlike in the expert mice (**c**), consistent with the population average (**f**). **i.** Comparison of cue response between trials with different lick times (calculated based on the grand average PSTH shown in **b** and **f**). *P*-value; hierarchical bootstrap testing the significance of correlation coefficient between lick time vs. cue-response (with a null hypothesis that there is no correlation). Error bar, SEM (hierarchical bootstrap). Left, schema showing time windows used to calculate cue response and pre-lick activity. SR, spike rate. Tcue, time from the cue. Tlick, time from lick. **j.** Same as in **i** for pre-lick activity. **k.** The time between 10% to 90% activity is calculated for the grand average PSTH shown in **b** and **f** (Methods). *P*-value; hierarchical bootstrap testing the significance of correlation coefficient between lick time vs. the time between 10-90% activity. Error bar, SEM (hierarchical bootstrap).

First, we leveraged trial-to-trial variability of lick time (CV = 0.41 ± 0.04; mean ± SEM; Extended Data Fig. 1k) to characterize how ALM preparatory dynamics change as a function of lick time in expert mice. Many ALM neurons showed ramping spiking dynamics starting at the cue onset and reaching a peak around the lick onset^2,52^ (Fig. 3a, cell 175, and Fig. 3b). When expert mice licked at different timing, the slope of ramping activity was altered without a change in the peak activity level (Fig 3a and b; trials with different lick times are shown in different colors). Consistently, temporal warping of spiking dynamics to normalize the lick time resulted in near-identical activity patterns across trials at both single-cell and population levels (even for non-ramping up cells, e.g., cell 232 in Fig. 3a; Fig 3b and c and Extended Data Fig. 5; Methods). Thus, the ALM preparatory dynamics are temporally stretched or compressed to trigger licking at different times in expert animals, consistent with previous observations in other tasks^51–53,63,64^.

Next, we examined whether similar reconfiguration in ALM dynamics underlies learning. At the beginning of delay training, mice licked early, and ALM neurons showed a transient response to the cue (Fig. 3e and f and Extended Data Fig.5; blue traces). Since strong ALM activity drives a lick^5,45^, this high amplitude cue response may explain immediate licks following the cue.

As mice licked later during learning, ALM dynamics progressively changed (Fig. 3e-f; from blue to red traces). First, the spiking activity following the cue decreased (‘cue response’; Fig. 3i), which may explain the loss of immediate licks. In contrast, the activity around the lick onset (‘pre-lick activity’) stayed high across trials (Fig. 3j), which may function as a ‘threshold’ activity level to trigger lick^51^. Second, slow dynamics (persistent or ramping activity) emerged, and filled the extended temporal gap between the cue and the delayed lick (Fig. 3k and Extended Data Fig. 7; the time between 10% to 90% activity level increased). These two modes of reconfigurations continued throughout delay training, converting the transient cue response into a ramping activity observed in the expert mice (Fig. 3f). Unlike in experts, ALM dynamics during learning were distinct across lick timings after the temporal warping (Fig. 3e-g).

During learning, Pearson’s correlation of ALM population activity between trials with different lick times is high across time points after the cue (Fig. 3h). This implies similar population activity patterns constitute preparatory activity across lick times (while the amplitude of activity changes). Consistently, a large proportion of the ALM preparatory activity is explained along a single dimension (65-80%; Extended Data Fig. 8), implying that learning reconfigures dynamics in a low-dimensional space^65^. In contrast, in the expert mice, population activity patterns changed between cue to lick, and thus the dimensionality is higher (Fig. 3d and Extended Data Fig. 8). Altogether, the reconfiguration of ALM preparatory dynamics during learning is qualitatively different from that in expert animals.

### Blocking CaMKII activity in ALM PT neurons impedes the evolution of dynamics

We next asked how the CaMKII manipulation in PT neurons influences the reconfigurations of ALM preparatory dynamics. To this end, we performed extracellular electrophysiological recordings in ALM in conjunction with PT neuron-specific paAIP2 manipulation during delay training (Fig. 4a).

**Figure 4.**
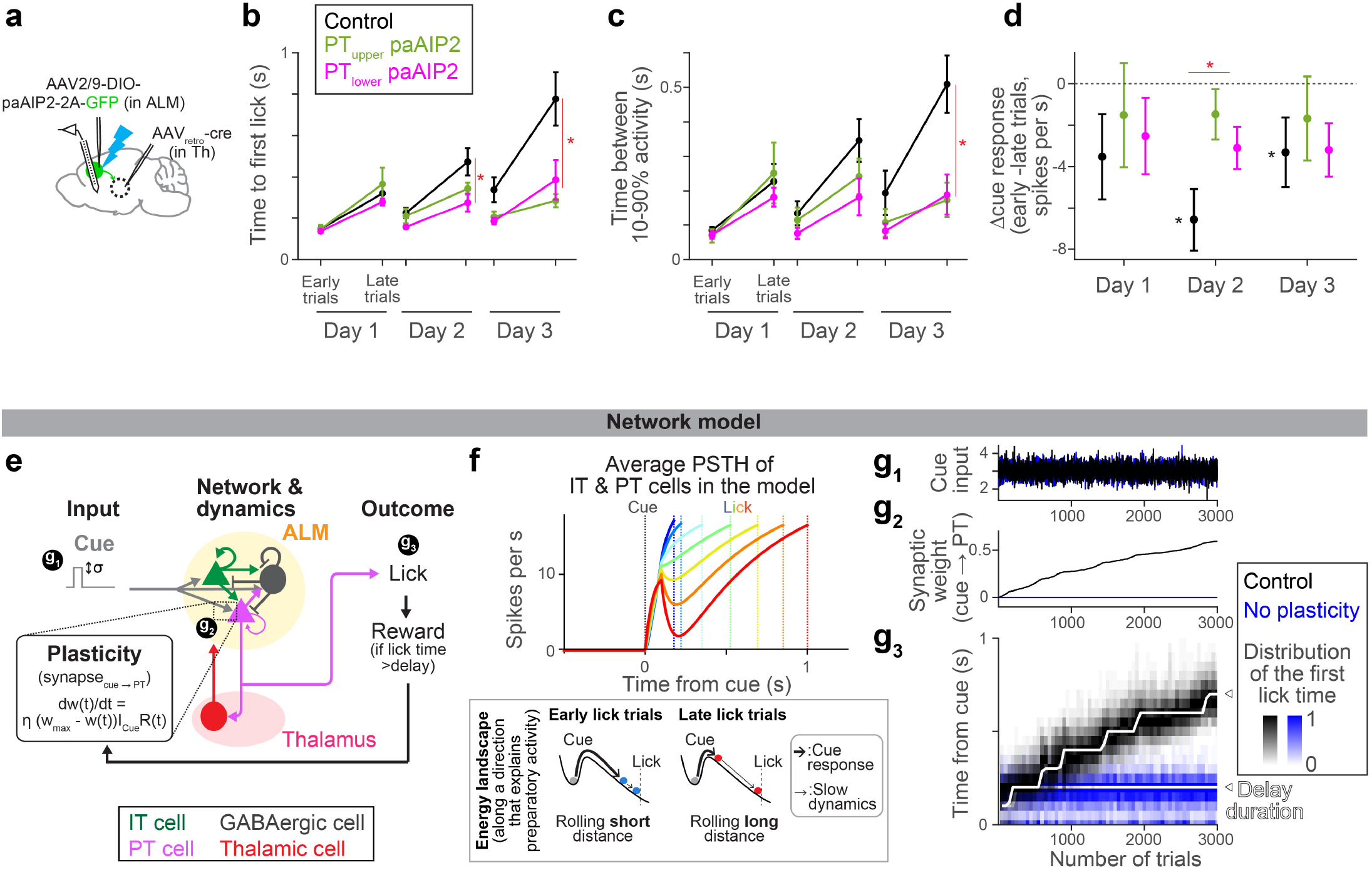
CaMKII manipulation in ALM PT neurons impedes the evolution of preparatory activity. **a.** Schema. Recording of extracellular activity in ALM during paAIP2 manipulation of PTupper (or PTlower) neurons. **b.** Time to first lick during delay training with recording (different cohorts of mice from those tested without recording in Fig.2c). Lines, mean ± SEM. *: *p* = 0.476, 0.005, <0.001 for day 1, 2, and 3 of delay training, respectively, comparing control vs. PT-specific paAIP2 manipulation (both PT cell types were pooled for statistics because we did not observe a qualitative difference between these two groups; the same in c and d; hierarchical bootstrap with a null hypothesis that the increase in lick time within a session is not larger in control). See Extended Data Table 3 for comparisons of each PT cell type and sample size. Early and late trials, first 75 and last 75 trials in the session. **c.** The time between 10% to 90% activity of positively-modulated ALM neurons is compared across manipulation types. Lines, mean ± SEM. *: *p* = 0.400, 0.095, 0.009 for day 1, 2, and 3, respectively, comparing control vs. PT-specific paAIP2 manipulation (both PT cell types were pooled; hierarchical bootstrap with a null hypothesis that the increase within a session is not larger in control). See Extended Data Table 3 for comparisons of each PT cell type and sample size. **d.** The change in cue amplitude of positively-modulated ALM neurons within sessions (late trials – early trials; Δcue response) of. Lines, mean ± SEM. * in red: *p* = 0.273, 0.006, 0.362 for day 1, 2, and 3, respectively, comparing control vs. PT-specific paAIP2 manipulation (both PT cell types were pooled; hierarchical bootstrap with a null hypothesis that the decrease within a session is not larger in control). * in black: *p* = 0.039, <0.001, 0.002 on day 1, 2, and 3, respectively, for hierarchical bootstrap with a null hypothesis that Δcue response is non-negative in control mice. **e.** Schema of the model (see Methods for details). All synaptic connections are excitatory, except for the ones from GABAergic neurons. “t” in the learning rule, trial. **f.** Top, dynamics of ALM neurons across lick times in the model (the mean of IT and PT neurons activity). Different color indicates activity in trials with different lick times. Dotted lines, corresponding lick times. Bottom, schemas of the energy landscape in early and late lick trials (along a long-time constant dimension that captures preparatory activity). Note that the full dynamics (top) is not monotonic due to activity along other directions. **g.** The amplitude of cue input (g1), synaptic weight of cue to PT synapse (g2), and lick timing (g3) during learning in the model.

To quantify how ALM dynamics change over training, we analyzed activity chronologically (Extended Data Fig. 9a). In control mice, the dynamics slowed down over three days of delay training, consistent with the delayed lick time (Fig.4b and c, black). In addition, the cue response significantly decreased within a session on days 2 and 3 of delay training (Fig. 4d; note that the cue response reaches the ‘floor’ as mice lick later, Fig. 3i, which may explain a stronger reduction on day 2). These reconfigurations of preparatory dynamics were significantly attenuated in the animals with PT neuron-specific paAIP2 manipulations (Fig. 4c and d and Extended Data Fig. 9a). Since preparatory activity precedes movement, the impedance in its reconfiguration likely explains the lack of learning during the paAIP2 manipulation.

Lick time and ALM cue response varied across trials, even during paAIP2 manipulations (Extended Data Fig. 7 and 9). In rare trials in which mice happened to lick late during PT neuron-specific paAIP2 manipulation, ALM dynamics were similar to that in the control (Extended Data Fig. 7-9). This implies that paAIP2 manipulation in PT neurons does not perturb spontaneous fluctuations in ALM dynamics. Instead, it is required to directionally reconfigure dynamics for learning.

### Potential mechanisms for synaptic plasticity to change lick time

How can synaptic plasticity in PT neurons reconfigure preparatory dynamics and drive learning? We generated a network model of ALM (Fig. 4e and Extended Data Table 4; Methods) constrained by previous findings: IT neurons form strong connections within the neocortex^66^. PT neurons, in contrast, do not project back to IT neurons but project to the thalamus and brainstem^66^ (for simplicity, we combined PT subtypes in the network model). The thalamic nuclei receiving PT input project back to ALM, and this thalamocortical loop maintains the preparatory activity^67,68^. The projection of PT neurons to the brainstem drives a lick when activity is high^45,47,48^.

This model implemented preparatory dynamics as ALM activity transitioning from the baseline activity to the threshold activity level that triggers a lick^2^ (Fig. 4f). At the beginning of a trial, the transient cue input rapidly pushes the activity (ball) out of the ‘baseline’ stable point, corresponding to the transient cue response (thick arrows, Fig. 4f bottom). After this transient excursion, the ball slowly rolls down the energy landscape until it reaches the threshold activity level to trigger a lick, which explains the slow ramping dynamics (thin arrows, Fig. 4f bottom). Thus, the amplitude of the cue response influences the following ramping dynamics (the energy landscape illustrates activity along a single dimension and does not explain the full dynamics, e.g., non-monotonic transient changes are affected by other dimensions). We varied the amplitude of the cue input across trials (Fig. 4e and g_1_), which changed the preparatory activity, and thus, induced across-trial variability in lick times as in the data. This allowed exploration of lick times required for learning^58^.

By imposing 1) a simple reward-dependent plasticity rule^69^ in the excitatory synapse between cue input and PT neurons and 2) the delay training protocol we applied to mice, the model reproduced the neural dynamics and learning observed in mice (Fig. 4e-g). A reward was provided when the network happened to ‘lick’ later than the delay. The reward-dependent plasticity potentiated the synapse, which paradoxically reduced the cue response due to strong excitatory inputs from thalamus/ALM neurons to the GABAergic neuron^70^ (alternatively, a model with synaptic depression without paradoxical effect reproduced similar dynamics and learning; Extended Data Fig. 10c-e). Consequently, in the following trials, the cue response was reduced, and the lick time was delayed. Thus, the plasticity allowed the network to exploit lick time guided by reward, and iterations of this cycle resulted in a gradual and directional change in preparatory dynamics and lick time (Fig. 4f and g). Without the synaptic plasticity in PT neurons, the model displayed no learning without a change in the variability of lick time, reproducing experimental observations (Fig. 4g, blue). The same network could reproduce expert dynamics by varying tonic input^71^ (Extended Data Fig. 10). Altogether, the model replicates the key experimental observations in this paper and proposes a potential mechanistic link among them: how synaptic plasticity in PT neurons shapes neocortical dynamics and behavior, i.e., decreased ALM cue response (Fig. 4d), generated slow dynamics (Fig. 4c), and delayed lick time (Fig. 4b).

## Discussion

We provided lines of evidence supporting synaptic plasticity in PT neurons as a learning mechanism in the motor timing task. First, we performed three types of acute cell-type-specific genetic manipulations targeting proteins required for synaptic plasticity, all of which blocked delay learning when PT but not IT neurons were manipulated. Acute manipulations are unlikely to recruit developmental effects or compensatory mechanisms^72^. Importantly, these manipulations did not affect the execution of learned behavior, implying that the effect is specific to learning^39,73^. Second, we performed a series of electrophysiological recordings *in* and *ex vivo* (Extended Data Fig. 2), confirming that transient paAIP2 manipulation does not affect the excitability of neurons or ongoing spiking activity. Instead, the paAIP2 manipulation impedes the directional reconfiguration of ALM dynamics, leading to rewarded lick, without perturbing trial-to-trial fluctuations in dynamics and behavior (Fig. 4 and Extended Data Fig.7-9). This is consistent with a view that CaMKII activity in PT neurons consolidates dynamics followed by a reward, presumably via synaptic plasticity of inputs that led to successful actions (Fig 4, model).

Synaptic plasticity has been observed across nearly all brain areas and cell types, including putative IT neurons^12,19,21^, seemingly suggesting global and redundant learning mechanisms. Thus, it was unexpected for a subset of cell types in one cortical area to be necessary for learning.

PT neurons occupy a unique position in the neocortex: they integrate the cortical input and control the information going through the thalamocortical loop that mediates preparatory activity^47,67^. Plasticity in PT neurons is, thus, well suited to reconfigure preparatory activity. Interestingly, putative PT neurons (complex-tuft cells) show active structural synaptic plasticity across areas and conditions^19,20,74^. In addition, localized synaptic plasticity, especially at the output nodes of a network like PT neurons, is theoretically beneficial^75,76^. Thus, PT neurons could be a universal learning site across neocortical areas and behavioral tasks. PT neurons encode behavior more reliably than other cortical cell types^46^, which could result from behaviorally relevant synaptic plasticity in these neurons.

Nonetheless, the necessity of one cell type does not exclude the necessity of other cell types and brain areas. Indeed, we discovered that at least two PT subtypes are required for learning. Neural dynamics are orchestrated across brain areas, including the striatum and cerebellum. Diverse cell types across these areas may implement different aspects of motor learning^23,73,77–79^. Molecular pathways underlying synaptic plasticity could also be diverse, and we do not rule out plasticity without CaMKII and Cofilin. Leveraging genetic manipulations targeting various molecules underlying synaptic plasticity, it is tempting to test how multiregional cell types coordinate diverse types of learning.

A common approach in neuroscience is to manipulate spiking activity and test its effect on behavior. Yet, spiking activity inevitably propagates to connected brain areas and often interferes with movement. For instance, silencing spikes in ALM alters spiking activity across motor-related brain areas and blocks licks^48,80^, which complicates the interpretation of behavioral effects even if it blocks learning^81^. The paAIP2 manipulations, instead, do not directly affect the ongoing spiking activity or execution of learned behavior. Molecularly-based plasticity manipulations have been used to test the causality of brain areas for learning^11,37,40,41,78,82^. We further advanced such methods with cell-type-specific manipulation combined with *in vivo* high-density electrophysiology during learning. This approach could effectively map learning mechanisms across behavioral tasks and brain areas.

Animals learn appropriate types and timing of movement. CaMKII manipulations in ALM blocked learning of lick timing, without affecting the acquisition of tongue movement to the lick port or cue association (Fig.1). Thus, different mechanisms may underlie learning movement types. Interestingly, learning movement types, such as sensory-motor association and learning new kinematics, generates new M1 activity patterns during movement across days^3–8,13^. Although we observed low-dimensional reconfiguration of ALM preparatory dynamics within sessions (Fig. 3), longitudinal recordings are required to test the emergence of activity patterns over days. In addition, our manipulations do not distinguish the type of synaptic plasticity responsible for learning. Furthermore, the pathways providing reward information and learning rule in PT neurons are unknown. The long-term goal is to explore these questions by directly and longitudinally monitoring synaptic plasticity and spiking activity of PT neurons during learning.

Impairments in learning and memory have devastating consequences in life. PT neurons are the homologous population of neurons vulnerable to amyotrophic lateral sclerosis and frontotemporal disorder in humans^83^. Identifying neocortical cell types required for learning, our findings and approach could be a foundation for future translational research of various diseases and injuries that affect learning.

## Acknowledgment

We thank N. Li, L. Colgan, T. Wang, D. Fitzpatrick, and Inagaki lab members for comments on the manuscript, M. Inagaki and L. Walendy for animal training, E.Wu, V. Grimaldi and R. DiCicco for histology, P. Scarpinato for DeepLabCut analysis, J. Yu for molecular biology, and H. Shearin and other MPFI ARC members for animal care. This work was funded by ZIA MH002497-34 (C.R.G.), Howard Hughes Medical Institute (S.R.), Max Planck Florida Institute for Neuroscience (H.K.I. and R.Y.), Max Planck Free Floater Program (H.K.I.), NIH New Innovator Award (NINDS and OD; 1DP2NS132108; H.K.I.), Searle Scholars Program (H.K.I.), Klingenstein-Simons Fellowship (H.K.I.), and McKnight Scholar Award (H.K.I.).

## Author Contributions

S.M., K.H., and H.K.I. planned and performed experiments and analyzed data. Z.Y., S.M., and H.K.I. developed the motor timing task. R.P., C.R.G. and K.H. performed histology. F.L. and S.R. performed network modeling. A.J. and R.Y. performed ex vivo recordings. S.M. and H.K.I. wrote the paper with input from all the authors.

## Author Information

The authors declare no competing interests. Correspondence and requests for materials should be addressed to.

## METHOD DETAILS

### EXPERIMENTAL MODEL AND SUBJECT DETAILS

#### Mice

This study is based on both male and female mice (age > P60, except for acute slice recordings that were P28). We used seven mouse lines: C57BL/6J (JAX# 000664), Tlx-Cre PL56 (RRID: MMRRC_041158-UCD)^61^, GRP-Cre KH288 (RRID: MMRRC_031183-UCD)^61^, Sim1-cre KJ18 (MGI: 4367070)^61^, LSL-Cas9 (B6J.129(B6N)-Gt(ROSA)26Sortm1(CAG-cas9-EGFP)Fezh/J, JAX# 026175)^62^, CaMKIIα-cKO (JAX# 006575)^60^ and, Vgat-ChR2-EYFP (JAX# 014548)^84^. Transgenic mice with non-C57BL/6J backgrounds were backcrossed to C57BL/6J for at least three generations. See Supplementary Table 1 for mice used in each experiment.

All procedures were in accordance with protocols approved by the MPFI IACUC committee. We followed the published water restriction protocol^85^ (with a modification of a minimum of 0.6 ml water per day to compensate for the high humidity in Florida). Mice were housed in a 12:12 reverse light: dark cycle and behaviorally tested during the dark phase. A typical behavioral session lasted between 1 and 2 hours. Mice obtained all of their water in the behavior apparatus. Mice were implanted with a titanium headpost for head fixation and single-housed.

#### Virus injection

We followed published protocols (dx.doi.org/10.17504/protocols.io.bctxiwpn) for virus injection. See Supplementary Table 1 for detailed descriptions of viruses and injection coordinates. See Supplementary Table 2 for a list of viruses used in this research^37,38,86^.

#### Behavior

A day before the training of the motor-timing task, we habituated mice with the experimental setup. A water-restricted mouse was head-fixed and placed in a training rig. 100 water drops (approximately 2 µL/drop) were delivered through a lickport at random timing (inter-trial intervals (ITI) sampled from an exponential distribution with a mean of 7.5s) without any sensory cue.

Next, we performed two-step training of the motor-timing task: cue association (2-3 days) and delay training (∼6 days). Both training phases shared the following general task structure: At the beginning of each trial, an auditory cue was presented, which consisted of three repeats of pure tones (3 kHz, 150 ms duration with 100 ms inter-tone intervals). A delay epoch started from the onset of the cue presentation. Licking during the delay epoch aborted the trial without water reward, followed by a 1.5 s timeout epoch. Licking during the 3 s answer epoch following the delay was considered a ‘correct lick’, and a water reward (approximately 2 µL/drop) was delivered, followed by a 1.5s consumption epoch. Trials without lick during the delay and answer epochs were considered ‘no response’ trials. Trials were separated by ITI randomly sampled from an exponential distribution with a mean of 2.2 s with 0.3 s offset (with a maximum ITI of 5 s). This prevented mice from predicting the trial onset without cue. Animals had to withhold licks during the full ITI epoch for the next trial to begin (if they licked, ITI was repeated). In a fraction of randomly interleaved trials, the auditory cue and water reward were omitted to assess spontaneous lick rate (‘no cue’ trials).

The cue association phase lasted for 2 or 3 sessions with 250-300 trials each. 15% of trials were no cue trials. The delay epoch duration was set to a minimum (0.1 s). After the second session, an animal was considered to have learned the cue association if the median lick time (the first lick time from the cue onset) in the last 100 trials was lower than 0.5 s. The few mice that failed to learn cue association within 3 sessions were excluded from the subsequent experiments (6/198 mice).

The cue association was followed by delay training which lasted for at least 6 days or until the animal reached a 1.8 s delay duration (beyond this delay, lick timing became too variable and unstable with our training condition). Sessions were terminated if animals stopped licking for 50 consecutive cue trials or reached 1000 trials. 5% of trials were no cue trials. The delay duration started at 0.1 s and automatically increased based on the performance: if the probability of correct trials in the last 100 cued trials with the same delay duration exceeded 30%, the delay duration was increased by 0.1s. The delay duration at the beginning of each session was set to 0.1s less than the final delay duration in the previous session (with a minimum delay duration of 0.1 s), as animals did not reach the criteria performance for the final delay duration on the last day.

For the electrophysiological recording of expert mice (Fig. 3 and Extended Data Fig.2g-o), animals were first trained up to 1.8 s delay following the protocol above. Then, the animals were trained with a 1.5 s delay duration for at least two weeks until their performance was stable with a masking flash (see ***Optogenetics***). The delay duration was fixed at 1.5 s during the recording.

To avoid bias, 1) behavior was automatically controlled by Bpod (Sanworks) and custom MATLAB codes, 2) experimenters were blinded to the genotype of mice, and 3) control and experimental animals were run in parallel.

#### Optogenetics

For paAIP2 experiments (Fig. 1 and 2c), animals were implanted with a clear skull cap^85^. Stainless steel tubes (0.062” OD, 0.052” ID, 8988K36, McMaster CARR) were cut into 3mm long pieces and glued directly on top of the clear skull cap above the target region (ALM or M1) to serve as sleeves for optic fibers. 470 nm LED light (M470F3, Thorlabs) at 0.5 mW power (power measured at the fiber tip) was delivered bilaterally through fiber optics (NA 0.39, 400 μm core diameter, M98L01, Thorlabs). The light was on for 1 second every 5 seconds (0.2 Hz) throughout the training session (LED illumination timing was independent of behavior). For control light-off sessions, the LED light was directed away from the skull with the same illumination protocol (i.e., mice could see flickering light as in the light-on condition).

For SN-CALI experiments (Fig. 2f), the same clear skull protocol was used. 595 nm LED light (M595F2, Thorlabs) at 0.75 mW power was delivered bilaterally for 1 minute every 10 minutes through optic fibers.

For paAIP2 manipulation during electrophysiological recordings (Fig. 4), 470nm LED light was delivered through an optic probe holder (OFPH_100_500-0.63_FC, Doric lenses) and probe tip (0.63NA, 500 μm core diameter, OPT_500-0.63_FLT, Doric lenses). The location of the optic probe tips was adjusted to cover the whole craniotomy (∼ 1.5 mm). 470 nm light at 3 mW power (at the probe tip) was delivered bilaterally for 1 second every 5 seconds (0.2 Hz).

For paAIP2 manipulation in expert mice (Fig.1e and Extended Data Fig. 2g-o), the light was turned on at least 30 minutes after the session onset. To prevent mice from distinguishing photostimulation and control trials, a ‘masking flash’ (1 ms pulses at 10 Hz) was delivered near the eyes using 470 nm LEDs (Luxeon Star) throughout the session.

#### Extracellular electrophysiology

A small craniotomy (diameter, 1-1.5 mm) was made over the recording sites a day before the first recording session. Extracellular spikes were recorded using 64 ch two-shank silicon probes (H-2, Cambridge Neurotech). Voltage signals were multiplexed, recorded on a PCI6133 board (National instrument), and digitized at 400 kHz (14-bit). The signals were demultiplexed into 64 voltage traces sampled at 25 kHz and stored for offline analysis. All recordings were made with the open-source software SpikeGLX (http://billkarsh.github.io/SpikeGLX/). During recordings, the craniotomy was immersed in a cortex buffer (125 mM NaCl, 5 mM KCl, 10 mM glucose, 10 mM HEPES, 2 mM MgSO_4_, 2 mM CaCl_2_; adjust pH to 7.4). Brain tissue was allowed to settle for at least five minutes before recordings.

#### Histology

Mice were perfused transcardially with PBS, followed by 4% PFA / 0.1 M PBS. Brains were post-fixed overnight and transferred to 20 or 30 % sucrose PB before sectioning on a freezing microtome. Coronal 50 µm free-floating sections were processed using standard fluorescent immunohistochemical techniques.

For images in Extended Data Fig. 4, all sections were stained with NeuroTrace® 435/455 Blue Fluorescent Nissl Stain (Thermo Fisher Scientific, N21479). The fluorescent label was amplified with chicken anti-GFP (Thermo Fisher Scientific, A10262) and goat anti-chicken 488 (Thermo Fisher Scientific, A11039). Slide-mounted sections were imaged on a Zeiss microscope with a Ludl motorized stage controlled with Neurolucida software (MBF Bioscience, Williston VT). Imaging was done with a 10× objective and a Hamamatsu Orca Flash 4 camera. Every section from the frontal pole through the brainstem were imaged.

For the CaMKII⍺ staining (Extended Data Fig. 3), GFP signal was amplified with rabbit anti-GFP (Thermo Fisher Scientific, A11122, RRID: AB_221569, 1:500) and goat anti-rabbit 488 secondary antibodies (Thermo Fisher Scientific, A11008, RRID: AB_143165, 1:500). The CaMKII⍺ was labeled with anti-CaMKII-α (6G9) (Cell Signaling Technology, #50049, RRID: AB_2721906, 1:250) and goat anti-rabbit 647 secondary antibodies (Thermo Fisher Scientific, A21236, RRID: AB_2535805, 1:500). Sections were imaged with a confocal laser scanning microscope (ZEISS, LSM 980) using a 20× objective.

#### Acute slice recording

AAV was injected into P28 mice to label PT_upper_ neurons in ALM bilaterally (n = 5; see Extended Data Table 1 for injection coordinate). Two to three weeks later, blue light (0.5mW 0.2Hz for one hour) illumination was given to one hemisphere under isoflurane sedation. Immediately after this, a slice was prepared and recorded (we also performed ex vivo illumination of brain slices, which yielded similar results; data not shown). Mice were perfused intracardially with a chilled choline chloride solution (124 mM Choline Chloride, 2.5 mM KCl, 26 mM NaHCO_3_, 3.3 mM MgCl_2_, 1.2 mM NaH_2_PO_4_, 10 mM Glucose, and 0.5 mM CaCl_2_, pH 7.4 equilibrated with 95 % O_2_ / 5 % CO_2_). The brain was removed and placed in the choline chloride solution. Transverse slices (300 μm) from both hemispheres containing the ALM were cut using a vibratome (Leica) and maintained in a submerged chamber at 32 °C for 1 h and then at room temperature in oxygenated artificial cerebrospinal fluid (ACSF: 127 mM NaCl, 2.5 mM KCl, 2 mM CaCl_2_, 1mM MgCl_2_, 25 mM NaHCO_3_, 1.25 mM NaH_2_PO_4_ and 25 mM glucose).

paAIP2-GFP labeled ALM PT_upper_ neurons were visualized using epifluorescent illumination. Whole-cell current-clamp recordings were obtained in labeled neurons using a Multiclamp 700B amplifier. Patch pipettes (3-6 ΩM) were filled with a potassium gluconate solution (130 mM K gluconate, 10 mM Na phosphocreatine, 4 mM MgCl_2_, 4 mM NaATP, 0.3 mM MgGTP, 3 mM L-Ascorbic acid, 10 mM HEPES. pH 7.2, 320 mOsm). These experiments were performed at room temperature (∼25° C) in oxygenated ACSF. Recordings were digitized at 10 kHz and filtered at 2 kHz. Current injections were given in 100 pA increments from −100 to 400 pA. Threshold, AP half-width, and time of AP half-width were analyzed in the current step where the first AP was observed. Recordings were performed in both hemispheres (one with illumination before recording and the other without), and the experimenter was blinded to the identity of the illuminated hemisphere during recording. All data were acquired and analyzed with custom-written C# and MATLAB codes.

#### Molecular Biology

For the cell-type-specific paAIP2 manipulation, we generated AAV-CaMKII-DIO-mEGFP-P2A-paAIP2. We used the same promoter (CaMKII promoter), 3’ sequence (bGH without WPRE sequence), and serotype (AAV9) with AAV-CaMKII-mGFP-P2A-paAIP2 (addgene: 91718), which we used for bulk manipulation (Fig. 1c), to match the expression level. The paAIP2 expression with CAG promoter and WPRE sequence in PT neurons, but not in IT neurons, resulted in light-independent blocking of learning, most likely due to over-expression (data not shown). We subcloned the mEGFP-P2A-paAIP2 sequence from the pAAV-CaMKII-mGFP-P2A-paAIP2 into the pAAV-CaMKII-FLEX-MCS plasmid (MPFI molecular core) using AscI and BamHI. For CRISPR/Cas9 KO of CaMKII⍺, we generated pAAV-U6CaMKIIgRNA-hsyn-mScarlet. We synthesized U6-CaMKII⍺ gRNA sequence^30^ and inserted it into pAAV-hSyn-mScarlet (addgene: 131001) using XbaI and ApaI. AAV2/9 based on these plasmids were packaged by UNC vector core.

For validation of efficacy of gRNA (Extended Data Fig. 3a and b), we transfected Neuro-2A cells in 6 well plates with pCAG-cre, pLenti-Cas9-GFP (addgene: 86145), and pAAV-U6CaMKIIgRNA-hsyn-mScarlet using Lipofectamine^TM^ 2000 (Invitrogen). Two days after lipofection, GFP+ mScarlet+ cells were sorted (BD FACSAria™ Fusion), and genome DNA was extracted using DNeasy Blood and Tissue Kit (QIAGEN). Index was added using PCR with Nextera XT Index Kit (Illumina). We used MiSeq 300 cycle ver. 2 (Illumina) for the sequencing.

### QUANTIFICATION AND STATISTICAL ANALYSIS

#### Behavioral analysis

Mice often ignored several cue trials at the beginning of each session. In addition, sated mice stopped licking at the end of sessions. To analyze behavior while mice are engaged in the task, we analyzed all trials between the first occurrence of 5 consecutive cue trials with licks and 20 trials before the last occurrence of 3 consecutive no-response trials.

We analyzed the time of the first lick after the cue (referred to as ‘lick time’). Electrical lick ports measured lick time, detecting the tongue’s contact with the lick port. To plot learning curves in Fig.1b and c (inset), all cue trials within each session were binned every 50 trials. Then, histograms of lick time were computed for each bin and were normalized to the respective peaks.

To compare learning across conditions (Figs.1, 2 and Extended Data Fig. 1 and 7), we used the last 100 cue trials in each session to compute the median first lick time, coefficient of variation (standard deviation divided by the mean) of the first lick time, and no response rate. To calculate lick time, ‘no-response’ trials were excluded. If mice reached a 1.8 s delay before 6 days of the delay training, the training was stopped, and the lick timing on the last behavior session was duplicated for analysis.

To compare the effect of paAIP2 manipulation in expert mice, we analyzed 3 blocks of trials for within-animal comparisons (Fig.1e, and Extended Data Fig.2k and l). In Fig. 1e, we had two control conditions: ‘Light off’, data of expert mice without paAIP2 manipulation (light onset was artificially and randomly assigned from those in Light on sessions); ‘Light on, shuffle’, data of light on sessions, but the light onset was randomly reassigned across sessions.

#### Videography analysis

High-speed (300 Hz) videography of orofacial movement (side view) was acquired using a CMOS camera (Chameleon3 CM3-U3-13Y3M-CS, FLIR) with IR illumination (940nm LED). We used DeepLabCut^87^ to track the movement of the tongue and jaw. Movements along the dorsoventral direction were analyzed and plotted in Extended Data Fig 1. Trajectories were normalized: the mean position before the cue was subtracted from trajectories and divided by the minimum value (thus, the downward movement of the tongue and jaw is upward in the plot). The onset of jaw movement in each trial is the first time point after the cue when the normalized movement trajectory exceeds 0.15. The onset of tongue movement is when DeepLabCut first detects the tongue after the cue.

#### Extracellular recording analysis

JRClust^88^ (https://github.com/JaneliaSciComp/JRCLUST) with manual curations was used for spike sorting. We used a combination of quality metrics to rigorously select single units for analysis (Extended Data Fig. 6). First, a false positive rate was estimated according to the inter-spike interval violation^89^, and units with a false positive rate < 0.15 were selected. Second, we extracted spike features (2 principal components of spike shape recorded in 3 neighboring recording sites) of all spikes recorded at the same or adjacent recording sites. Then, we calculated the Mahalanobis distance of spike features of each spike from the center of the spike cluster (unit) of interest. We performed the receiver operating characteristic (ROC) analysis of Mahalanobis distance to distinguish spikes within vs. outside the cluster of interest. A recording session was divided into time bins containing consecutive 1000 spikes of the unit of interest. AUC was calculated for every time bin. Units with a mean AUC (across time bins) > 0.9 and without any time bin with AUC<0.75 were selected. Third, units with low spike rates (less than 0.1 spikes per s) were excluded from the analysis. Units passing all these criteria were deemed single units.

We recorded 3689 neurons passing the quality metrics in ALM from 30 mice during learning. Putative pyramidal neurons (units with spike width > 0.5 ms ^90^; 3203 among 3689 neurons) were analyzed. To define preparatory cells, we compared the spike rates in the baseline (0 to 0.2 s before the cue) vs. during the task (from cue to first lick) in trial 21-95 of the session (‘Early trials’ in Fig. 4a). Cells with a *p*-value lower than 0.5 (signed-rank test) were considered task-modulated cells and analyzed in Figs. 3 and 4 (See Extended Data Table 3 for details). Cells with positively and negatively task-modulated cells (cells with significant increase or decrease in spike rate during the task) were analyzed separately (except for Fig.3 c, d, g, and h) so that they do not cancel out. Since there were more positively modulated cells (Extended Data Fig. 7e), the main figures show the analysis of positively modulated cells.

For the peri-stimulus time histograms (PSTHs) in Fig. 3 and Extended Data Fig. 5, trials were pooled based on the first lick time. Analyzed time ranges were T = [0.05∼0.20, 0.20∼0.35, 0.35∼0.50, 0.50∼0.65, 0.65∼0.80, 0.80∼0.95, 0.95∼1.1] second for recording during learning, and T = [0.95∼1.1, 1.1∼1.25, 1.25∼1.4, 1.4∼1.55, 1.55∼1.7, 1.7∼1.85, 1.85∼2.0] second for recording in expert mice. Ten trials with the first lick within the time range were randomly selected, and the spike rate was averaged. If the number of trials within the range was less than 10, the range was not included in the analysis. PSTHs were smoothed with a 50 ms causal boxcar filter. For Extended Data Fig. 2h, PSTHs were smoothed with a 20 ms causal boxcar filter to detect fast changes in activity. SEM was based on hierarchical bootstrap: first, we randomly selected animals with replacement; second, we randomly selected sessions of each animal with replacement; and third, we randomly selected cells within each session with replacement (1000 iterations).

To temporally warp the PSTH (Fig. 3), we linearly scaled the spike timing in the time range T_cue_ > 0.1 by (LT_target_ - 0.1)/(LT_trial to be warped_ - 0.1), where T_cue_ denotes time after the cue, and LT denotes the first lick time in each trial type (LT_target_ was 1 for Fig.3a,b, and f). After the warping, we calculated PSTH and smoothed it with a 50 ms causal boxcar filter. We did not warp 0 <T_cue_ < 0.1 as the onset of cue response showed the same temporal profile regardless of the lick time (most likely determined by the latency of cue input to ALM^48^). Because of this, we did not warp the first trial type with a lick latency between 0.05∼0.20 s during learning.

Cue response (Fig. 3 and 4, Extended Data Fig.2n) was normalized for the baseline spike rate: ^&SR^, where SR_T_ denotes mean spike rate in the time range indicated in T. Similarly, pre-lick activity was normalized by the baseline spike rate. To calculate ‘Time between 10-90% activity’ (Fig. 3 and 4), PSTH was normalized by the minimum and maximum spike rate between cue and lick. Then, the time from the first time point crossing 10% to the last time point crossing 90% was measured (in the case of negative modulated cells, Extended Data Fig. 7, activity was flipped). Mean speed (Extended Data Fig. 7) is the mean of absolute change in spike rate between cue to lick (here, spike rate was calculated in 50 ms time bin). We performed hierarchical bootstrap for statistics and SEM (1000 iterations).

For the correlation analysis (Fig. 3d and h, and Extended Data Fig. 7g), we looked for an *n* × 1 unit vector *r_T, trial type_*, representing the spike rate of all *n* neurons at time point T of a trial type of interest. Then, we calculated Pearson’s correlation of *r* between the two trial types indicated in the plot across time points (in the case of Extended Data Fig. 7g, autocorrelation was calculated).

For recording in the expert mice (Extended Data Fig. 2), we recorded 716 neurons that passed the quality metrics in ALM in 20 sessions from 4 expert mice. 647 putative pyramidal neurons were analyzed.

#### Ramping direction analysis (Extended Data Fig. 8)

To calculate ramping mode (**RM**) for a population of *n* recorded neurons, we looked for an *n* × 1 unit vector that maximally distinguished the mean activity before the trial onset (0 - 0.2 s before cue; ***r***_before cue_) and the mean activity before the first lick (0 – 0.15 s before the first lick; ***r***_before lick_) in the *n-dimensional* activity space. We defined a population ramping vector: ***w*** = ***r***_before lick_ – ***r***_before cue_. **RM** is ***w*** normalized by its norm.

In each recording session, we randomly selected 24 cue trials with the first lick time above 0.25 s. We define RM based on the activity of these randomly selected trials, and projected activity in different sets of trials for the cross-validation. To calculate the “activity explained”, we calculated the squared sum of the spike rate after subtracting the baseline (mean spike rate 0 – 0.2 s before cue onset) across neurons. We calculated the squared sum of the activity along RM after subtracting the baseline.

#### Histology Analysis

For Extended Data Fig. 3e, cell counting and signal analysis were conducted using MATLAB. Cells were binary classified into GFP and/or RFP positive/negative based on the fluorescence signal intensity. The CaMKII⍺ signal intensity within the cell was normalized by the signal in the surrounding areas (median of areas within 12-60 pixels (0.6 µm/pixel) from the cell).

For Extended Data Fig. 4, imaged sections were processed with NeuroInfo software (MBF Bioscience, Williston, VT) to align the serial sections into a whole brain volume. Neurons throughout the ALM cortex were detected and marked using the automatic cell detection function in NeuroInfo software, which uses a modified Laplacian of Gaussian (LOG) algorithm that detects labeled neuron perikarya based on size and fluorescence intensity with a neural network that eliminates signal artifacts. The accuracy of cell counts compared to visual detection is approximately 95%.

#### Network Model

Using a dynamical systems approach^91,92^, we consider four variables representing the average membrane currents (ℎ) and spike rates (*r* = *f*(ℎ), where *f*(ℎ) is the neural activation function) of neuronal populations in different regions of the brain. Specifically, we modeled two excitatory populations (pyramidal tract, ℎ_PT_, and intratelencephalic, ℎ_IT_) and one inhibitory population (ℎ_IN_) in the premotor cortex, one excitatory population in the thalamic nuclei receiving PT input (ℎ_th_). The average membrane current dynamics of population *k* are described by the following nonlinear differential equation:

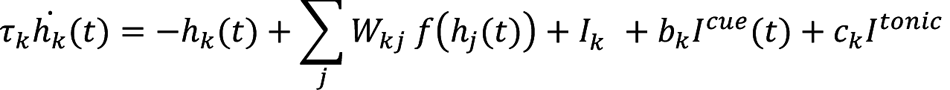

Where τ_k_ is the membrane time constant of population *k*, W_kj_ are elements of the connectivity matrix between presynaptic population *j* and postsynaptic population *k*, I^DC^_k_ is the baseline input current. I^cue^(*t*) is the external input current provided to ALM neurons elicited by the auditory cue via the synaptic weights *b*_k_, and I^tonic^ is the non-contextual tonic input provided to the thalamus via the synaptic weights c_k_. We hypothesized the pathway providing cue input is different from thalamic nuclei maintaining preparatory activity according to previous work^48^. For all populations, we used a threshold-linear activation function:

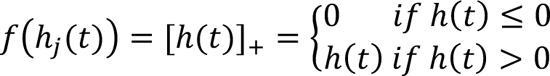

The baseline input currents were chosen to be negative for all four populations so that the network dynamics displays a baseline stable fixed point (lower attractor) for ℎ_5_(*t*) < 0 ∀ A *j*. The connectivity matrix **W** respected the biological constraints found in the literature (see Extended Data Table 4), and was chosen so that a stable fixed point (higher attractor) existed for ℎ_5_(*t*) > 0 ∀ *j*. The transition from the lower to the higher attractor was triggered by I^cue^(*t*) which took the form of a 100ms long boxcar function. Ramping activity emerged as the dynamics evolved towards the higher attractor following the cue. The lick onset was defined upon reaching a spike rate threshold (*r*^∗^_PT_ = 15 spikes per second) motivated by the activity of ALM (Fig.3).

We imposed synaptic plasticity between the input relaying the auditory cue and PT neurons (*b*_PT_) to implement learning. To update *b*_PT_, we used a heavily simplified learning algorithm. We hypothesized that the external inputs to the network I^cue^(*t*) and I^tonic^ were subject to fluctuations across trials. In particular, the peak value of the cue signal I^cue^ and I^tonic^ were each drawn from normal distributions with means μ^cue^, μ^tonic^ and standard deviations σ^cue^, σ^tonic^, respectively(see Extended Data Table 4). Because of these fluctuations, the spike rate of PT neurons will reach the threshold at a different speed in each trial. The synaptic weight *b*_0&_ was either exclusively potentiated (model shown in Fig.4), or depressed (model shown in Extended Data Fig. 10c-e), at the end of rewarded trials, i.e., trials in which the PT spike rate exceeded the hard threshold after the delay duration according to the update rule:

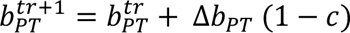

where Δb_PT_ = η [b^max^_PT_ − b^tr^_PT_]_+_I^cue^ for potentiation (Δb_PT_ = η [b^min^_PT_ − b^tr^_PT_]_+_I^cue^ for depression), η being the learning rate. *c* is the recent reward rate (mean of the last five trials; we added this term to prevent the network from learning when the reward rate was 100%). As in the behavioral experiments with mice, when the reward rate in the last 100 trials reached 30%, the delay duration was increased by 100 ms.

To model the behavior and the neural dynamics in expert mice (Extended Data Fig. 10a, b, and e), we used the same network parameters that were found at the end of the learning process. To vary the lick time, we changed the non-contextual tonic input I^3627’^(*t*), which is compatible with a previous paper modeling dynamics in expert animals performing a timing task^71^. In the synaptic potentiation model (Extended Data Fig. 10a-b), stronger tonic inputs lead to slower dynamics, allowing the model to capture the behavior of expert mice with increasing delay durations. In the synaptic depression model (Extended Data Fig. 10e), weaker tonic inputs lead to slower dynamics.

### Statistics

The sample sizes are similar to the sample sizes previously published in the field. No statistical methods were used to determine the sample size. During spike sorting, experimenters could not tell the trial type and therefore were blind to conditions. All *signed rank* and *ranksum* tests were two-sided. All bootstrapping was done at least over 1,000 iterations.

### Reagent and data availability

The new plasmids reported in this paper will be posted to Addgene before publication. The recording data in NWB format and example codes will be shared on DANDI around the time of publication.

---

**Extended Data Figure 1.**
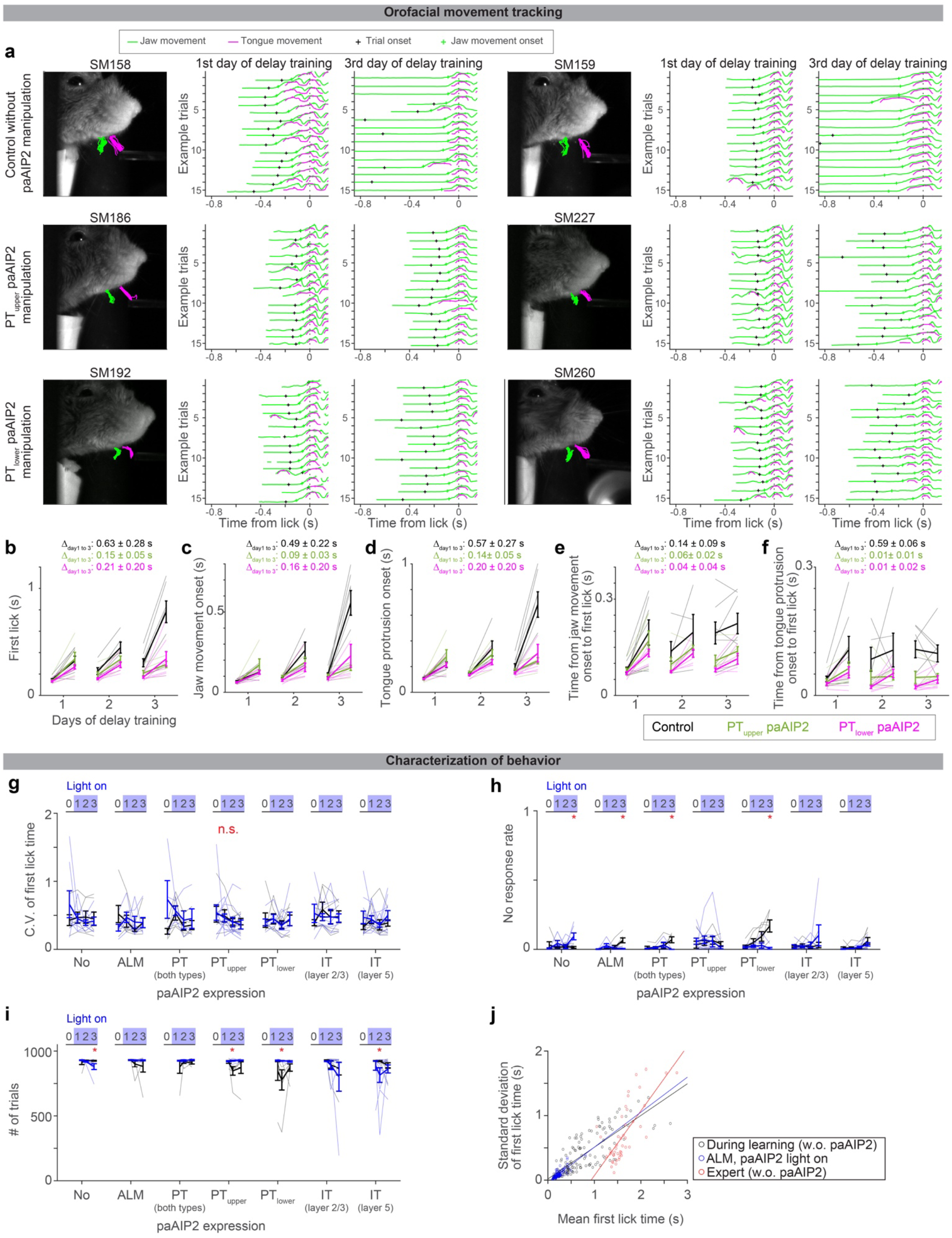
Characterization of behavior during delay training. **a.** Tracking of jaw and tongue, using high-speed videography and DeepLabCut^87^. Six example animals (2 per experimental condition) are shown. Left, video clip overlayed with example trajectories of jaw (green) and tongue (magenta) (15 trials). Middle, trajectories of jaw and tongue at the beginning of delay training (from trial 50 to 65 on day 1 of delay training). Trajectories are aligned to lick onset, where lick onset is defined as the moment tongue touches the lick port for the first time after the cue. Black cross, cue (trial) onset. Green cross, the onset of jaw movement (Methods; analyzed in panels **c** and **e**). Right, the same as in the middle but for later in delay training (trial 50 to 65 from the end of day 3 of delay training). **b-f.** Summary of the first lick time (**b**), jaw movement onset (**c**), tongue protrusion onset (**d**), time from jaw movement onset to the first lick (**e**), and time from tongue protrusion onset to the first lick (**f**) (Methods). Comparing the first and last 100 trials within a session. Thick lines, mean ± SEM. Thin lines, individual mice (n = 7, 5, and 7 for control, PT_upper_ paAIP2 manipulation, and PT_lower_ paAIP2 manipulation, respectively). The delayed tongue and jaw movement (i.e., withholding of orofacial movement after the cue onset) mainly explains the delayed lick time. **g-i.** Characterization of behavior during paAIP2 manipulation. The coefficient of variation (C.V.) of first lick time (**g**). No-response rate (**h**). Numbers of trials per session (**i**). These are based on animals and sessions analyzed in Fig. 2c. Thick lines, mean ± SEM. Thin lines, individual animals. No change in C.V. between control vs. paAIP2 manipulation indicates that the manipulation did not affect the variability of lick timing. No significant increase in the no-response rate and no significant decrease in the number of trials in paAIP2 manipulation implies that the manipulation did not block cue-triggered lick and motivation (note that all significant changes are in the opposite direction). *: *p* < 0.05 (bootstrap followed by *Bonferroni* correction; Extended Data Table 3); n.s.: *p* > 0.05 for all comparisons. **a. j.** Relationship between mean vs. standard deviation of first lick time. The standard deviation of lick time scales with its mean, adheres to Weber’s law^50,52^. This is why we analyzed CV (standard deviation divided by mean) to evaluate across-trial variability in lick time (e.g., **g**). Circle, the first 100 or last 100 trials of each session. Lines, least-square fitted lines. The slope of the fitted line: 0.49 (0.43 – 0.56), 0.53 (0.45-0.64), and 1.00 (0.77-1.35), mean (5 – 95 % confidence interval, bootstrap) for during learning (control), during learning (paAIP2 in ALM with light on), and expert, respectively. Thus, ALM paAIP2 manipulation does not affect the variability of lick time.

**Extended Data Figure 2.**
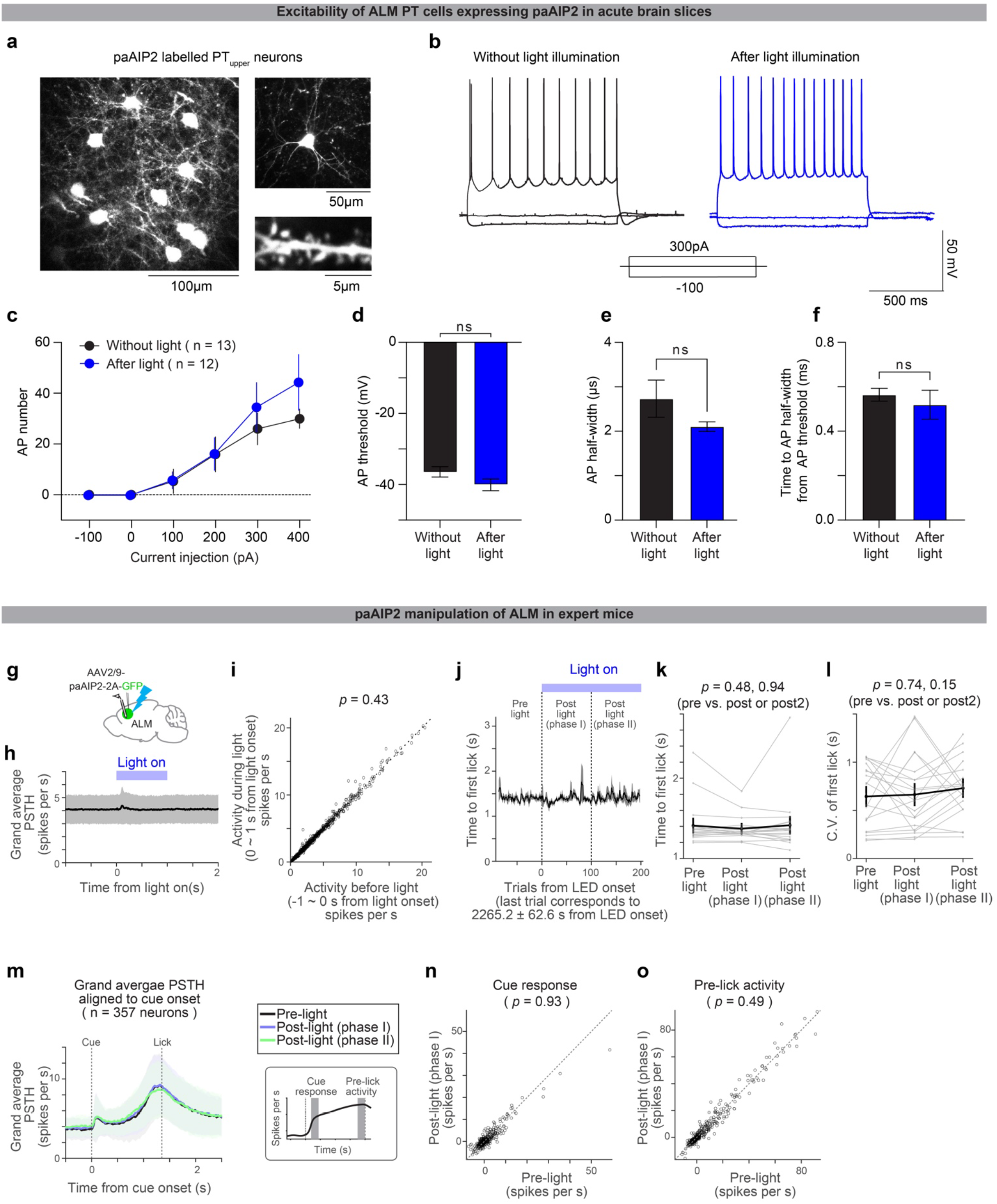
The paAIP2 manipulation does not affect excitability or ongoing spiking activity. **a.** Retrograde labeling of PT_upper_ neurons in ALM with GFP (co-expressed with paAIP2) in an acute cortical slice. Right top, example cell. Right bottom, example dendritic shaft. **b.** Representative traces of whole-cell current-clamp recordings from paAIP2 labeled PT_upper_ neurons at three different current steps (−100, 0, and 300 pA) without (black) and after (blue) 60 minutes of *in vivo* blue light stimulation (1 sec on, 4 sec off; Methods). **c.** The mean number of action potentials (AP) evoked by depolarizing current steps. n = 13 cells after stimulation and 12 cells before stimulation. Error bar, SEM. **d.** AP threshold showed no difference between control and stimulation (*p* = 0.1225, ranksum test). **e.** AP half-width showed no difference between control and stimulation (*p* = 0.3981, ranksum test). **f.** Time to the first AP half-width from AP threshold. There was no difference between control and stimulation (*p* = 0.7795, ranksum test). **g.** Schematic for the extracellular electrophysiological recording of ALM with paAIP2 manipulation in expert mice during behavior (delay duration, 1.5 s). Blue light (470 nm, 1 s on, 4 s off, 3 mW; Methods) was turned on >30 mins after the session onset. **h.** Blue light-triggered average of ALM activity. Thick line, average PSTH of all putative pyramidal ALM neurons (n = 647 neurons, 4 animals, 20 sessions). Blue bar, blue light pulse (1 s). Shade, SEM. We observed a non-significant increase in spike rate (∼0.2 spikes per s, on average) around 100ms after the light onset. This could be a visual response, as we observed a similar response in mice without paAIP2 expression (data not shown). **i.** Average spike rate of individual neurons before and during blue light pulses, calculated over 1s time windows before and during the blue light pulse. Circle, neuron. Dashed line, unit line. No difference in spike rate was observed before and during blue light (*p* = 0.69, signed rank test), indicating that paAIP2 manipulation does not directly affect spiking activity. **j.** Median lick time (5 trial sliding windows; aligned to the first trial with blue light illumination). Thick line, average across sessions (20 sessions from 4 animals). Shaded area, SEM. Vertical dashed lines separate 3 phases in the session: pre-light (0 - 100 trials before blue light onset), post-light phase I (0 - 100 trials after blue light onset), and post-light phase II (100 - 200 trials after blue light onset). The last trial corresponds to ∼40 mins of blue light illumination. **k.** Median lick times in the 3 phases in **j**. Thick line, mean ± SEM. Thin lines, individual sessions. No significant difference was observed (*p*-value, signed rank test), indicating that paAIP2 manipulation did not directly affect lick time in expert mice. **l.** Coefficient of variation of the first lick time in the 3 phases in **j**. The same format as in **k**. No significant difference was observed (*p*-value, signed rank test), indicating that paAIP2 manipulation does not directly affect the variance of lick time. The no-response rate did not change significantly, either (data not shown). **m.** Grand average PSTH aligned to the cue onset (for trials with lick times between 0.55 - 2.63 s, corresponding to 10% to 90% quantiles of the lick time distribution in these sessions). n = 357 significantly modulated neurons in ALM. Lines, mean. Shade, SEM. **n.** Comparisons of task-related activity (cue response: mean spike rate over 100-150 ms after cue onset minus baseline spike rate) before and after blue light illumination. Circle, neuron. Dashed line, unit line. We found no significant differences (signed rank test), suggesting that paAIP2 manipulation does not affect task-related activity in expert mice. **o.** The same as **n** but for pre-lick activity (mean spike rate over 50 ms preceding the first lick minus baseline spike rate). We found no significant differences (signed rank test), suggesting that paAIP2 manipulation does not affect task-related activity in expert mice.

**Extended Data Figure 3.**
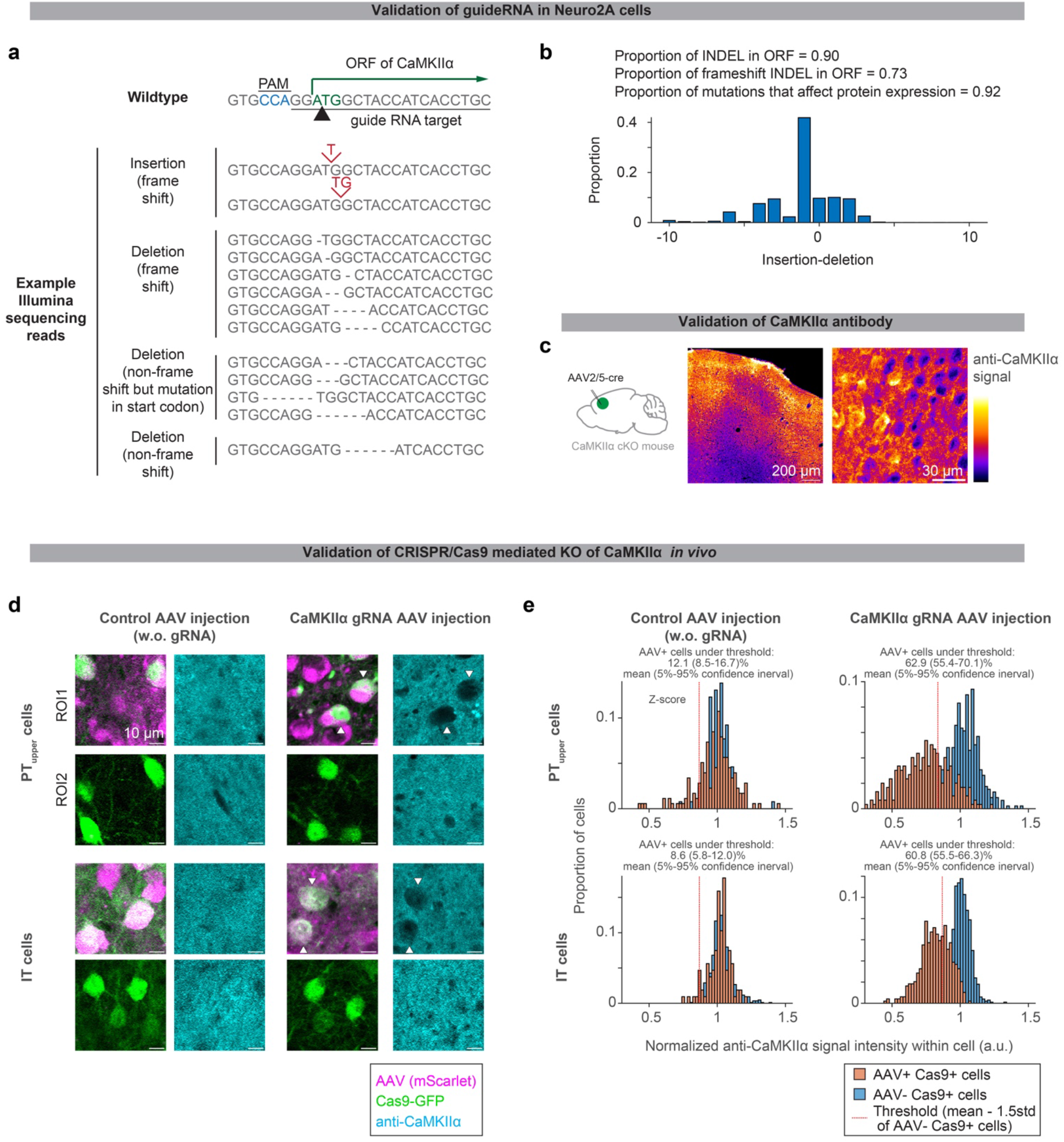
Histological validation of CRISPR/Cas9 KO of CaMKII⍺. **a.** Sequencing of genomic DNA in Neuro2A cells transfected with guide RNA against CaMKII⍺ and Cas9 (Methods). **b.** The proportion of amplicon with Indel mutation. **c.** CaMKII⍺ immunohistochemistry of ALM in CaMKII⍺ conditional knock out (cKO) mouse injected with AAV-hsyn-cre. Loss of signal was observed around the injection site (ALM; the center of the image), validating the CaMKII⍺ antibody. **d.** CaMKII⍺ immunohistochemistry of ALM with cell-type-specific CRISPR/Cas9 KO of CaMKII⍺. Loss of CaMKII⍺ was observed only in cells expressing both Cas9 (green) and guide RNA (magenta, in right panels). Black areas smaller than PT/IT neurons are presumably glia, inhibitory neurons, or neuropile of manipulated cells. **e.** Quantification of CaMKII⍺ immunostaining signal (Methods).

**Extended Data Figure 4.**
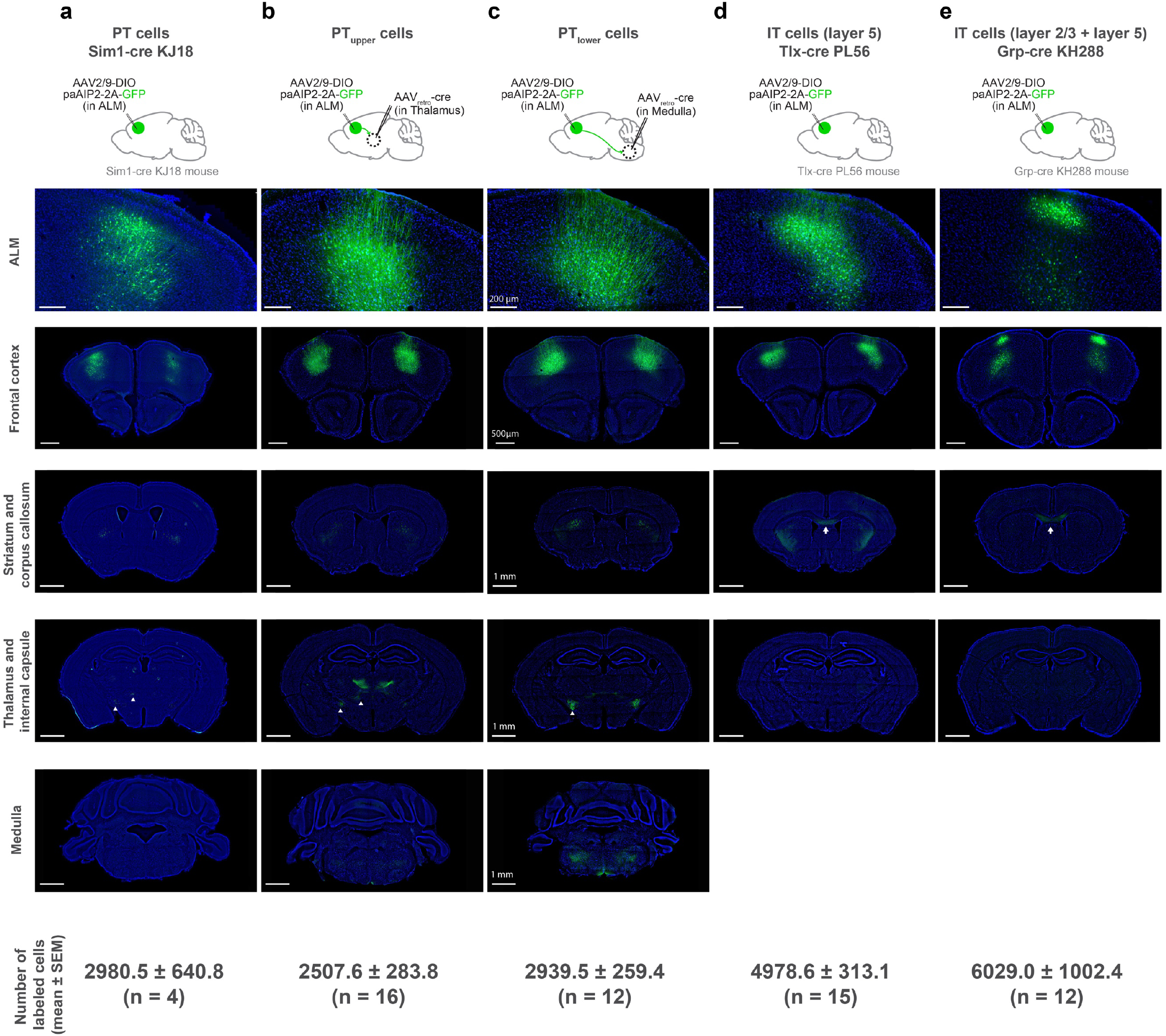
Histological validation of cell-type-specific manipulation. **a.** paAIP2-GFP expression in ALM PT neurons and some layer 2/3 neurons in Sim1-cre KJ18 mice. **b.** paAIP2-GFP expression in ALM PT_upper_ neurons (due to the tropism of AAV_retro_^47^, layer 6 neurons are only sparsely labeled). **c.** paAIP2-GFP expression in ALM PT_lower_ neurons. **d.** paAIP2-GFP expression in ALM layer 5 IT neurons in Tlx-cre PL56 mice. **e.** paAIP2-GFP expression in layer 2/3 IT and some layer 5 IT neurons in Grp-cre KH288 mice. Top, paAIP2-GFP expression in ALM. Bottom panels, paAIP2-GFP expression in 4 coronal sections from anterior to posterior showing expression in the frontal cortex, striatum and corpus callosum, thalamus, and medulla. See Extended Data Table 1 for injection coordinates. Signal in the corpus callosum (white arrows) in **d** and **e** confirms the labeling of IT neurons. Whereas the lack of signal in the corpus callosum in **b** and **c** confirms a lack of co-labeling of IT neurons. Signal in the thalamus and internal capsule (white arrowheads) in **a**, **b**, and **c** confirms the labeling of PT neurons. In contrast, the lack of signal in the thalamus and internal capsule in **d** and **e** confirms a lack of co-labeling of PT neurons. The labeling of PT neurons in Sim1-cre KJ18 mice (**a**) was sparser than the labeling of PT_upper_ or PT_lower_ neurons by AAV_retro_ (i.e., **b** and **c**). In addition, we note that Sim1-cre KJ18 labels some layer 2/3 neurons in ALM. Scale bars are identical across panels in each row.

**Extended Data Figure 5.**
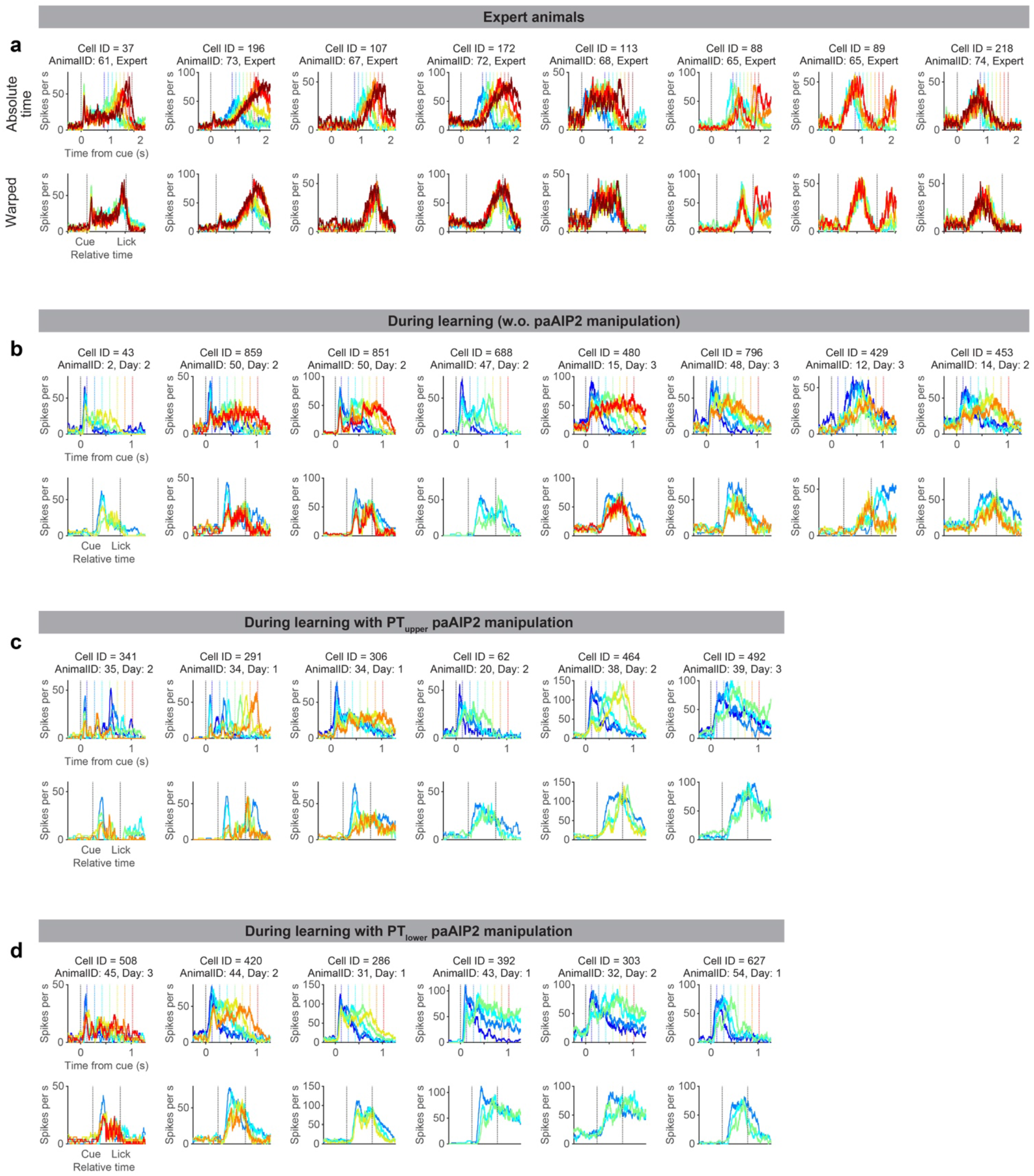
Spiking activity patterns of single cells Example neurons recorded during learning and in experts. The data format is the same as in **Fig. 3a**. Note that cells recorded in different animals often show similar reconfiguration of dynamics (e.g., Cell 43, 859, and 480 during learning without paAIP2 manipulation), indicating the reconfiguration is general across animals.

**Extended Data Figure 6.**
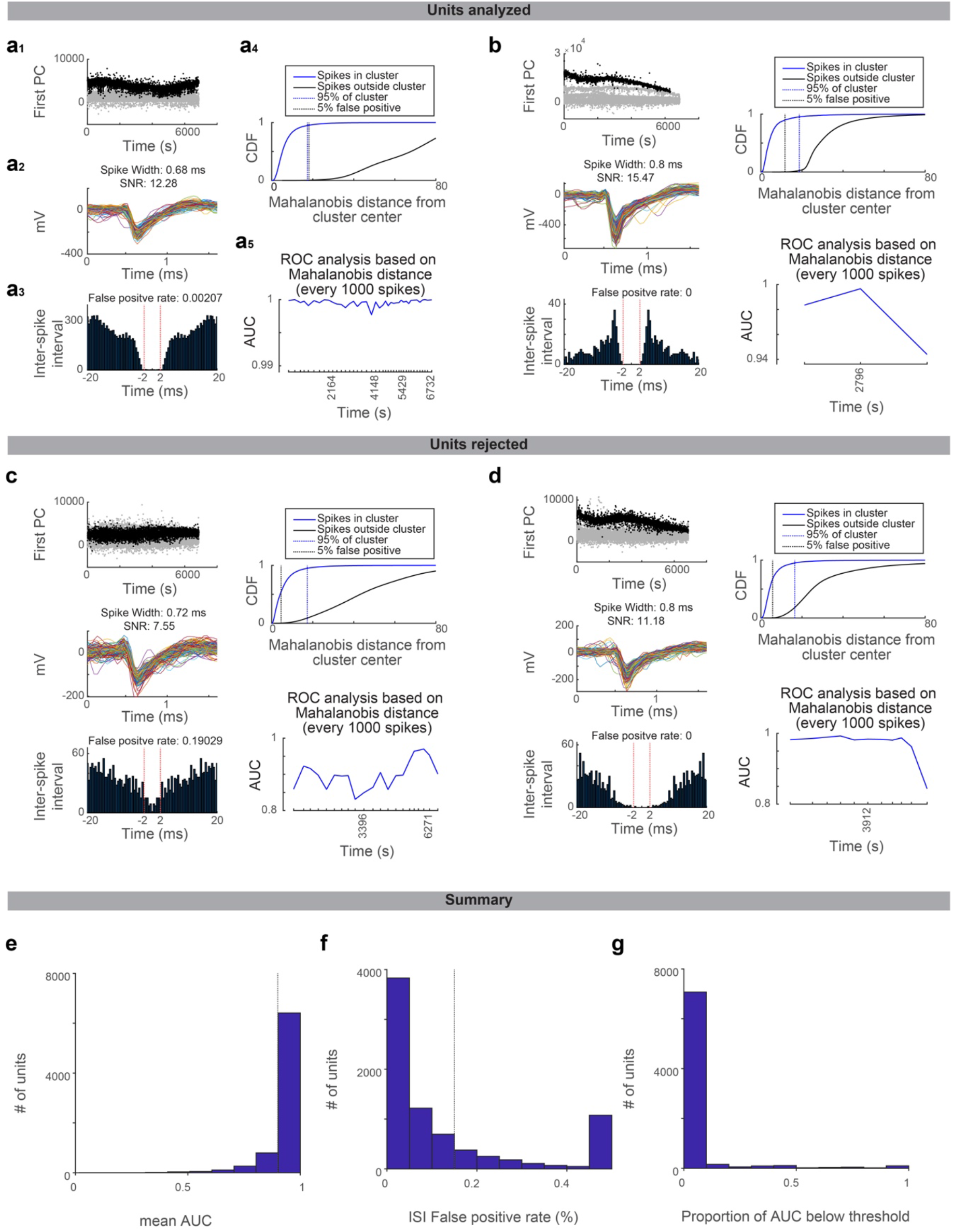
Quality metrics of spike-sorted units Drifts in the recording affect spike sorting quality. We implemented rigorous quality metrics evaluating cluster isolation across time points within a recording session to minimize the effect of recording drift in our analysis. **a.** An example unit that passed the quality standard. **a_1_**, projection of spike feature along the first PC. Time, the time within the recording session. Black dots, spikes belonging to the unit. Gray dots, all the other spikes sharing the peak channel (i.e., recorded at the same or adjacent recording sites). **a_2_**, spike shape of the unit belonging to this unit. Randomly selected 250 spikes are overlaid. **a_3_**, the inter-spike interval of spikes belonging to the unit (see Methods for false positive rate). **a_4_**, Mahalanobis distance (calculated based on spike features) of each spike from the center of the unit cluster. CDF is shown for spikes belonging to the unit (blue) and all other spikes sharing the peak channel (black). Blue dotted line, 95% point of spikes belonging to the unit. Black dotted line, the distance where the false positive rate reaches 5%. **a_5_**, ROC analysis of the Mahalanobis distance (distinguishing spikes belonging or not belonging to the unit; Methods). AUC is calculated for each time window containing every consecutive 1000 spikes belonging to the cluster. **b.** The same as in **a** for different unit **c-d.**The same as in **a** for different units not passing the quality standard due to high false alarm rate in inter-spike interval analysis (c) or mean AUC lower than the threshold (d). **e** The histogram of mean AUC (AUC averaged across all time windows) across all manually curated units. Quality criterion: mean AUC > 0.9 (dashed line). **f** The histogram of the false-positive rate calculated according to the inter-spike interval across all manually curated units. **g** No single time window should have AUC < 0.75 to pass the quality criteria. The proportion of AUC below the threshold (across all manually curated units) is shown for all manually curated units.

**Extended Data Figure 7.**
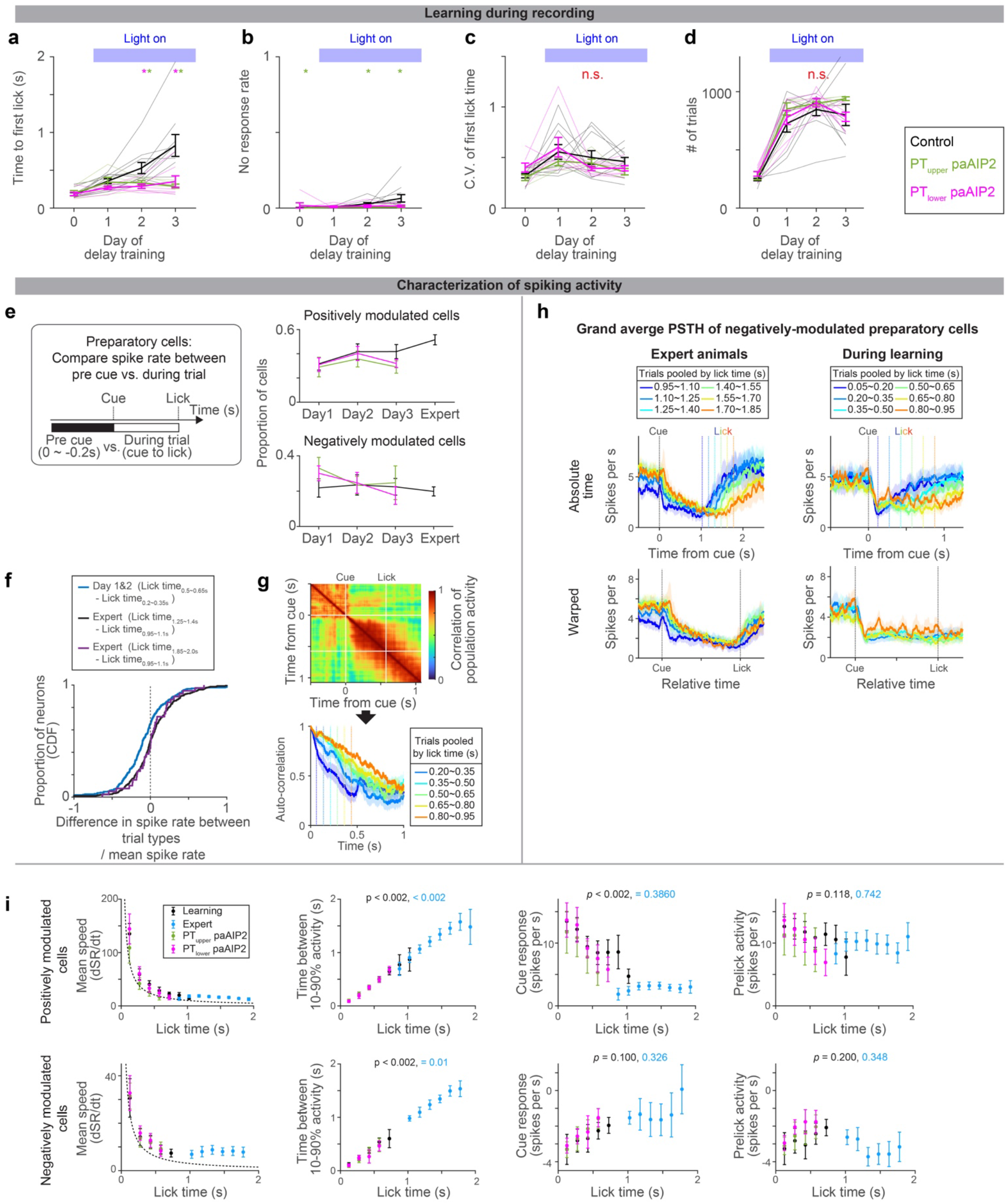
ALM dynamics during learning. **a-d.** Behavioral performance of the animals during ALM recording (different cohorts of mice from those analyzed in Fig2c; the same cohorts of mice analyzed in Fig.4). Time to first lick (**a**), no-response rate (**b**), coefficient of variation (**c**), and number of trials per session (**d**) are shown. Thick lines, mean ± SEM. Thin lines, animals. N = 10, 5, 7 mice for control, PT_upper_, and PT_lower_ manipulation, respectively. *: p< 0.05 control vs. PT_upper_ manipulation or control vs. PT_lower_ manipulation conditions (indicated by the color of *; bootstrap followed by *Bonferroni* correction; Extended Data Table 3), n.s., *p* >0.05 for all comparisons. The result is consistent with behavior without recording (Fig. 2 and Extended Data Fig. 1). **e.** The proportion of cells with positive or negative-modulated preparatory activity (spike rate between cue to the first lick significantly higher or lower from that in the baseline, repsectively; two-sided signed-rank test, *p* < 0.05). Error bar, SEM (hierarchical bootstrap). *p* > 0.05 for all comparisons of proportion (on days 1, 2, and 3) between control vs. manipulations (hierarchical bootstrap). Thus, the paAIP2 manipulation does not affect the proportion of preparatory neurons. **f.** Spike rate significantly decreased during learning. *(SR_trial type 1_ − SR_trial type 2_)/SR_both trial types_* was calculated for each neuron and the distributions of this value are shown as CDF, where ^SR^ denotes the mean spike rate between cue to first lick. During the training, a significant proportion of neurons decreased spiking activity as mice licked later (blue; *p* = 1.18 × 10^-6^, signed-rank test, n = 269 cells; median −0.103; based on neurons shown in Fig. 3g). In contrast, in the expert, there was no significant change in spike rate (black; *p* = 0.27, signed-rank test, n = 344 cells; median: −0.001; based on neurons shown in Fig. 3c) even when the fold-difference in lick time between trial types is roughly matched to that in Day1&2 (purple; *p* = 0.69, signed-rank test, n = 39 cells; median: 0.006). Altogether decrease in spike rate is unique to during learning. **g.** Autocorrelation of ALM population activity (top; trials with lick time between 0.50 and 0.65 s) to estimate time-constant of population activity (bottom; different trials are shown in different colors). The correlation of population activity between time points: (T_cue_ + T_lick_)/2 vs. following time points, is shown, where T_cue_ and T_lick_ denote the time of cue and lick, respectively. Vertical dotes lines, lick times. Consistent with **Fig. 3**, the time constant of population dynamics increased as mice learned to lick later. **h.** Grand average peri-stimulus time histogram (PSTH) of negatively-modulated ALM preparatory neurons. The same format as in **Fig. 3b**. Shade, SEM. Trial types with more than 25 neurons are shown. **i.** Characterization of PSTH of positively and negatively modulated ALM preparatory neurons. The format is the same as in **Fig. 3i-k**, and the data of paAIP2 manipulation is overlaid. Mean speed, the average spike rate (SR; spikes per sec) change between cue to lick. Dotted line, expected mean speed (mean pre-lick activity divided by lick time). The mean speed (first column) decreases, and time between 10-90% activity (second column) increases as animals lick later, consistent with a view that dynamics are temporally stretched. The absolute values of the cue response (third column) decrease during learning, whereas pre-lick activity (fourth column) is stable in both positively and negatively modulated cells. Trial types with more than 50 trials were analyzed.

**Extended Data Figure 8.**
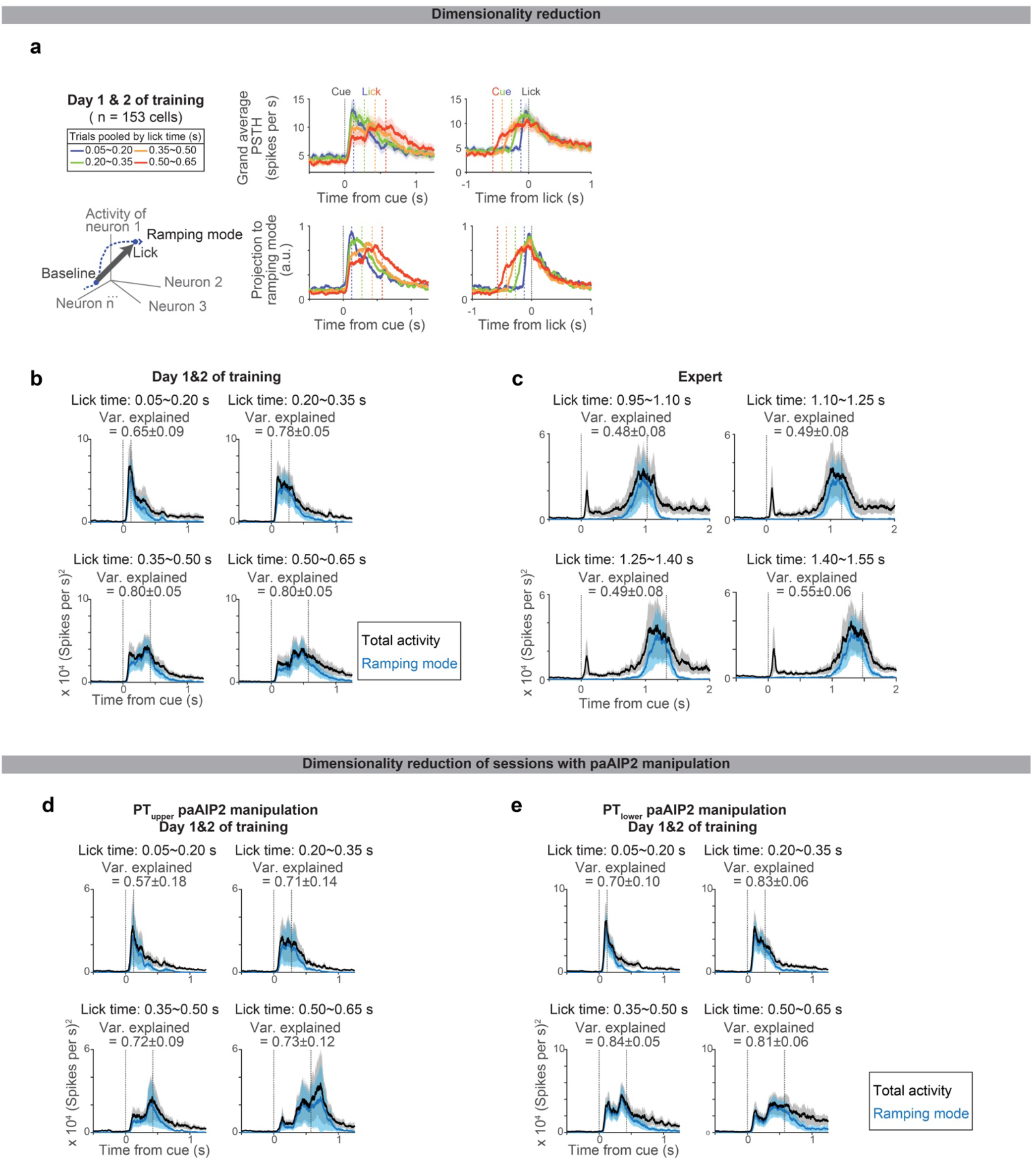
Low dimensional population activity during learning. **a.** Projection of ALM spiking activity to the ramping mode, which maximally distinguishes the population activity between the baseline and pre-lick (Methods). Top, grand average PSTH aligned to cue (left) or lick (right). Bottom, projection to the ramping mode. n = 153 preparatory neurons in ALM (neurons with >= 10 trials for all lick times were analyzed). **b.** The square sum of total spiking activity (black) and the square of projection along ramping mode (blue) at each time point, indicating the large proportion of spiking activity can be explained by activity along this mode. Shade, SEM (hierarchical bootstrap). Var explained: mean ± SEM. **c.** Same as **b** for Expert. N = 123 preparatory neurons in ALM (neurons with >=10 trials for all lick times were analyzed). **d.** Same as **b** for PT_upper_ paAIP2 manipulation, indicating the large proportion of spiking activity can be explained by activity along this mode even during the paAIP2 manipulation. N = 117 preparatory neurons in ALM (neurons with >=10 trials for all lick times were analyzed). **e.** Same as **b** for PT_lower_ paAIP2 manipulation. N = 140 preparatory neurons in ALM (neurons with >=10 trials for all lick times were analyzed).

**Extended Data Figure 9.**
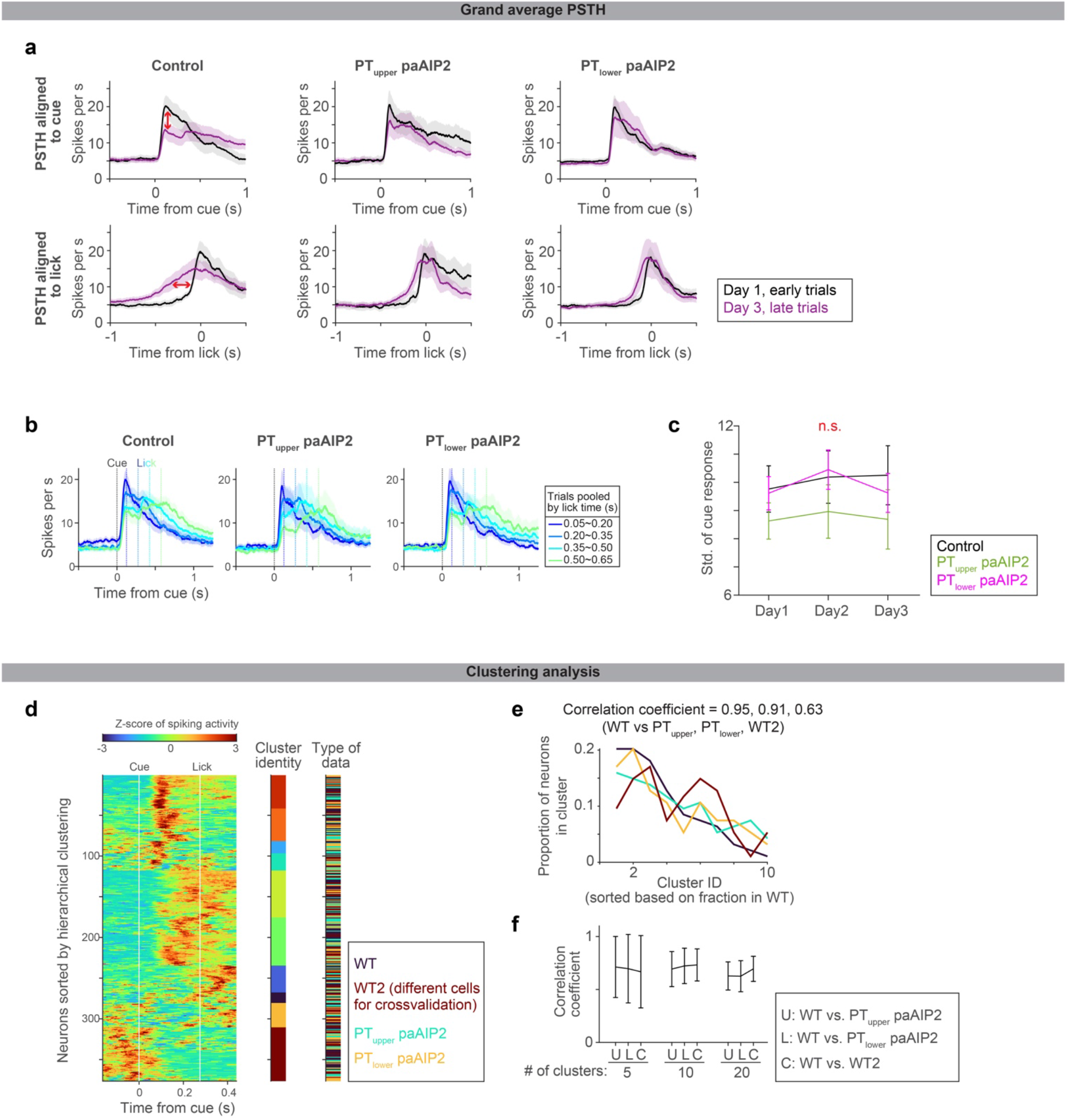
Similar ALM activity across experimental conditions when mice licked around the same time. **a.** Comparisons of the grand average PSTHs of positively-modulated ALM preparatory neurons on day 1 early trials vs. day 3 late trials (regardless of lick time; the same dataset is quantified in **Fig. 4c-d**). PSTHs are aligned to cue (top) or lick (bottom). Control shows a clear change in cue response and ramping activity (red arrows). Early and late trials, first 75 and last 75 trials in the session. See Extended Table 3 for number of neurons analyzed. **b.** When mice lick around the same time (regardless of the day of training), the grand average PSTHs of ALM are similar regardless of paAIP2 manipulation. The grand average PSTHs of positively-modulated ALM preparatory neurons for individual manipulation types are shown. Activity in trial types with different lick timings is shown in different colors. The paAIP2 manipulation sessions had fewer trials with later licks. The same format as in **Fig.3f**. Shade, SEM. The control data is duplicated from **Fig. 3f** for comparison purposes. **c.** The standard deviation of cue response (related to **Fig. 4d**; calculated based on early trials; late trials yield similar results). The standard deviation and CV (not shown) of cue response are not significantly different between control and paAIP2 manipulation. *p* = 0.284, 0.322, 0.274 (day 1, 2, 3 comparing control vs. PT_upper_ paAIP2 manipulation) and 0.830, 0.800, 0.592 (day 1, 2, 3 comparing control vs. PT_lower_ paAIP2 manipulation; hierarchical bootstrap). Although the across-trial fluctuation in cue response is the same, the mean cue response changed between early and lick trials only in control (**Fig. 4d**). Thus, the paAIP2 manipulation blocks the directional change in cue response without affecting the across-trial fluctuation. **d.** Clustering analysis to test whether we can distinguish ALM activity patterns between control and paAIP2 manipulation conditions. Z-scored spiking activity of ALM neurons recorded in control animals (WT) and animals with PT-specific paAIP2 manipulations is shown. Neurons from each experimental condition were subsampled and pooled (94 neurons per condition were randomly sampled without replacement; two different groups of neurons were randomly subsampled in the WT condition for cross-validation and denoted WT and WT2). Then we performed hierarchical clustering of the mean activity pattern of these neurons (here, the number of clusters was set to 10). Mean spiking activity in trials with a first lick time between 0.2 and 0.35 s was analyzed. The cluster identity of each neuron is shown in different colors in the middle column, and the type of experimental condition is shown in the right column (see legend in the figure for color scheme). Each cluster contains neurons recorded in all experimental conditions, implying that ALM neurons with similar activity patterns were recorded across experimental conditions. **e.** Fraction of neurons in each cluster in panel **d**. The cluster ID is sorted based on the proportion of neurons in WT data. The correlation coefficients of the proportion of neurons between experimental conditions, indicating similarity in the composition of activity patterns, are shown at the top. **f.** We have repeated this correlation analysis shown in **d-e** 1000 times (sampling random subsets of neurons without replacement; we have tested different numbers of clusters: 5, 10, or 20, all of which yielded similar results) to plot the mean and SEM. C, control: the correlation between WT vs. WT2, showing the upper bound of the correlation coefficient with this procedure and sample size (as neurons are sampled from the same dataset). Both PT_upper_ and PT_lower_ paAIP2 manipulation conditions have similar correlation coefficients to control (*p* > 0.05), indicating that activity patterns in ALM are indistinguishable across conditions when mice lick around the same time, i.e., paAIP2 manipulations in PT neurons do not change task-related activity patterns in ALM.

**Extended Data Figure 10.**
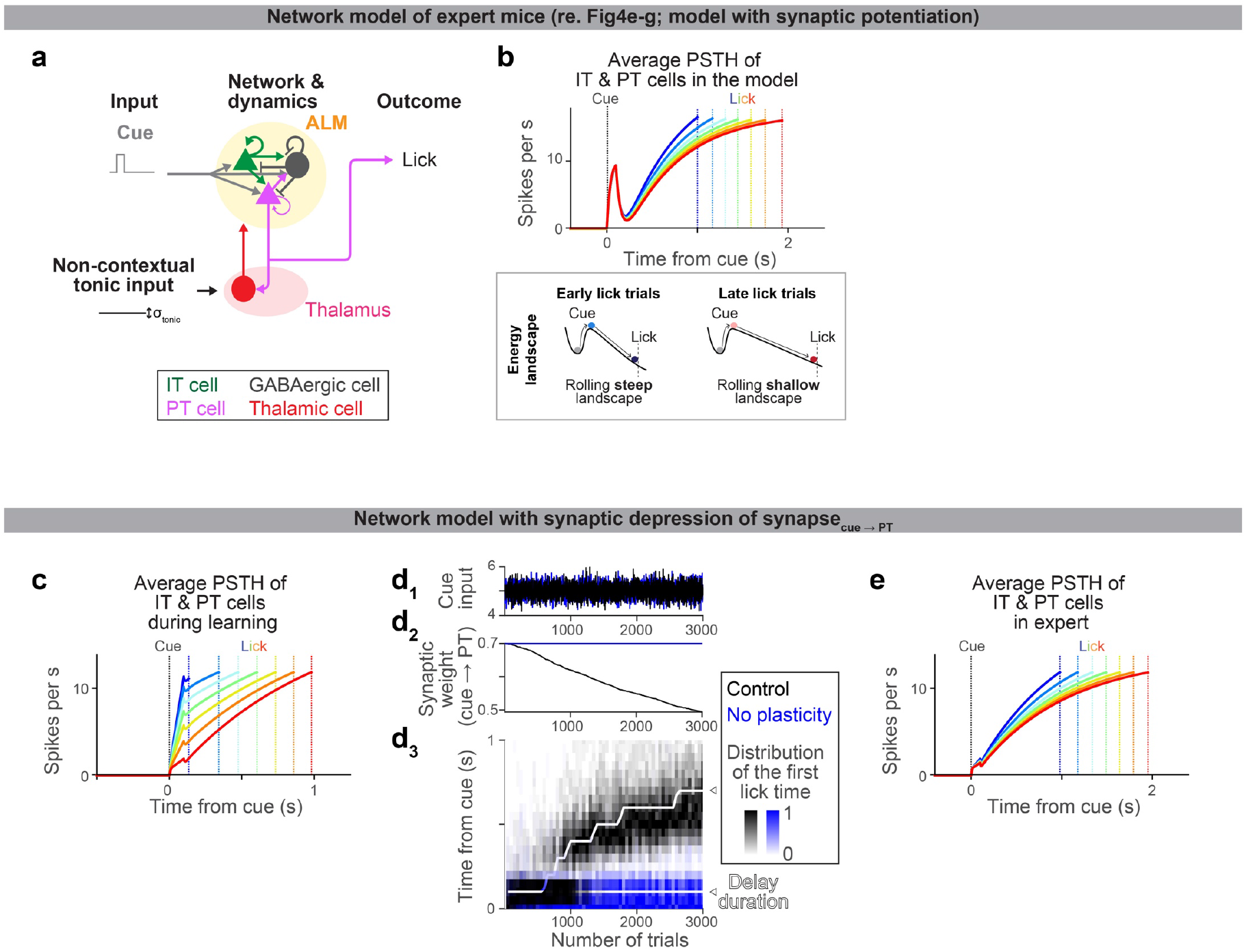
A model of expert dynamics and a model with synaptic depression. **a.** Schema of the expert model (see Methods for details). The network structure is identical to that for learning (**Fig. 4e**). Following literature^52^, we provided a non-contextual tonic input to vary lick times across trials (no plasticity is imposed as this is a model of expert). **b.** Dynamics of ALM neurons in the model (top) and corresponding energy landscape (bottom). Different color indicates activity in trials with different lick times. The amplitude of tonic input changed the slope of the landscape, which changed the speed of dynamics. This reproduced the ‘temporal scaling’ of ramping dynamics consistent with experimental data (**Fig.3**) and previous report^52^. **c.** The same format as in **Fig.4f**, but for a network model with synaptic depression of the synapse between cue and PT neurons. The network architecture is identical to that in **Fig.4e**, but with different synaptic weights (Extended Data Table. 4) and reward-dependent synaptic depression instead of potentiation (Methods). **d.** The same format as in **Fig.4g**, but for the network model with synaptic depression of the synapse between cue and PT neurons. **e.** The same format as in **b**, but for the network model with synaptic depression of the synapse between cue and PT neurons. Altogether, similar to the potentiation model, the depression model can reproduce the experimental observations.

