## Extended Data Table 1 for "Cell-type-specific plasticity shapes neocortical dynamics for motor learning"

Extended Data Table. 1: Details of surgery performed for each experiment in the paper.

| Figure | Purpose | Virus | Viral injection coordinates (all bilateral; unit 'mm') | Mouse line | Number of mice |
|---|---|---|---|---|---|
| Fig. 1c | No paAIP2 Control | N/A | N/A | C57BL/6J | 7 light on + 6 light off control |
| | paAIP2 manipulation in ALM | AAV2/9 CaMKII promoter-mEGFP-P2A-paAIP2 (v1) | Bregma AP 2.5, ML ±1.5, DV 0.4 and 0.8, 75 nl at each depth | C57BL/6J | 6 light on + 7 light off control |
| | paAIP2 manipulation in M1 | AAV2/9 CaMKII promoter-mEGFP-P2A-paAIP2 (v1) | Bregma AP 2.5, ML ±1.5, DV 0.4 and 0.8, 75 nl at each depth | C57BL/6J | 4 light on |
| Fig. 1d | paAIP2 manipulation in ALM during cue association | AAV2/9 CaMKII promoter-mEGFP-P2A-paAIP2 (v1) | Bregma AP 2.5, ML ±1.5, DV 0.4 and 0.8, 75 nl at each depth | C57BL/6J | 3 light on + 5 light off control |
| Fig. 1e | Expert control, light off | N/A | N/A | C57BL/6J or Vgat-ChR2-EYFP | 4 C57BL/6J + 6 Vgat-ChR2-EYFP, light off |
| | Expert with paAIP2 manipulation in ALM, light on | AAV2/9 CaMKII promoter-mEGFP-P2A-paAIP2 (v1) | Bregma AP 2.5, ML ±1.5, DV 0.4 and 0.8, 75 nl at each depth | C57BL/6J | 4 light on |
| Fig. 1f | Control for CaMKIIα conditional knockout | AAV2/5 hsyn promoter-cre (v4) | Bregma AP 2.5, ML ±1.5, DV 0.8, 110 nl | Wildtype littermates of CaMKIIα-cKO mice | 7 |
| | CaMKIIα conditional knockout in ALM | AAV2/5 hsyn promoter-cre (v4) | Bregma AP 2.5, ML ±1.5, DV 0.8, 110 nl | CaMKIIα-cKO flox/flox | 7 |
| Fig 2c | paAIP2 manipulation in ALM PT neurons | AAV2/9 CaMKII promoter-DIO-mEGFP-P2A-paAIP2 (v3) | Bregma AP 2.5 mm, ML ±1.5 mm, DV 0.8 mm , 110 nl | Sim1-cre KJ18 | 4 light on + 5 light off control |

| | | | | | |
|---|---|---|---|---|---|
| | paAIP2 manipulation in ALM $PT_{upper}$ neurons | AAVretro CamKII promoter-Cre, 50% dilution (v2) in thalamus | Bregma AP -1.5 mm, ML ±1.0 mm, DV 3.2 mm , 100 nl | C57BL/6J | 11 light on + 7 light off control |
| | | AAV2/9 CaMKII promoter-DIO-mEGFP-P2A-paAIP2 (v3) | Bregma AP 2.5 mm, ML ±1.5 mm, DV 0.4 mm and 0.8 mm, 75 nl at each depth (or 75 or 110 nl at DV 0.8 mm) | | |
| | paAIP2 manipulation in ALM $PT_{lower}$ neurons | AAVretro CamKII promoter-Cre, 50% dilution (v2) in Medulla | Bregma AP -6.65 mm, ML ±1.25 mm, DV 4.5 mm , 200 nl | C57BL/6J | 6 light on + 7 light off control |
| | | AAV2/9 CaMKII promoter-DIO-mEGFP-P2A-paAIP2 (v3) | Bregma AP 2.5 mm, ML ±1.5 mm, DV 0.4 mm and 0.8 mm, 75 nl at each depth (or 75 or 110 nl at DV 0.8 mm) | | |
| | paAIP2 manipulation in ALM layer 5 IT cells | AAV2/9 CaMKII promoter-DIO-mEGFP-P2A-paAIP2 (v3) | Bregma AP 2.5 mm, ML ±1.5 mm, DV 0.8 mm , 110 nl | TLx-Cre PL56 | 10 light on + 8 light off control |
| | paAIP2 manipulation in ALM layer 2/3 IT cells | AAV2/9 CaMKII promoter-DIO-mEGFP-P2A-paAIP2 (v3) | Bregma AP 2.5 mm, ML ±1.5 mm, DV 0.8 mm , 110 nl | GRP-Cre KH288 | 6 light on + 6 light off control |
| Fig. 2e | CRISPR/Cas9 KO of CaMKII α in ALM PT cells | AAVretro CamKII promoter-Cre, 50% dilution (v2) in thalamus | Bregma AP -1.5 mm, ML ±1.0 mm, DV 3.2 mm , 100 nl | LSL-Cas9 | 9 |
| | | AAV2/9 U6 promoter CaMKII gRNA-hsyn promoter-mScarlet (v5) | Bregma AP 2.5 mm, ML ±1.5 mm, DV 0.8 mm , 110 nl | | |

| | | | | | |
|---|---|---|---|---|---|
| | CRISPR Control in ALM PT cells | AAVretro CamKII promoter-Cre, 50% dilution (v2) in thalamus | Bregma AP -1.5 mm, ML ±1.0 mm, DV 3.2 mm , 100 nl | LSL-Cas9 | 8 |
| | | AAV2/1 hSyn promoter-mScarlet (v6) | Bregma AP 2.5 mm, ML ±1.5 mm, DV 0.8 mm , 110 nl | | |
| | CRISPR/Cas9 KO of CaMKII α in ALM IT cells | AAV2/9 U6 promoter CaMKII gRNA-hsyn promoter-mScarlet (v5) | Bregma AP 2.5 mm, ML ±1.5 mm, DV 0.8 mm , 110 nl | TLx-Cre PL56 x LSL-Cas9 | 6 |
| | CRISPR Control in ALM IT cells | AAV2/1 hSyn promoter-mScarlet (v6) | Bregma AP 2.5 mm, ML ±1.5 mm, DV 0.8 mm , 110 nl | TLx-Cre PL56 x LSL-Cas9 | 6 |
| Fig. 2f | Cofilin-SuperNova in ALM $PT_{upper}$ cells | AAVretro CamKII promoter-Cre, 50% dilution (v2) in thalamus | Bregma AP -1.5 mm, ML ±1.0 mm, DV 3.2 mm , 100 nl | C57BL/6J | 6 light on + 7 light off control |
| | | AAV2/9 Ef1a promoter-DIO-CFL-SN (v7) | Bregma AP 2.5 mm, ML ±1.5 mm, DV 0.8 mm , 110 nl | | |
| | Supernova-Control in ALM $PT_{upper}$ cells | AAVretro CamKII promoter-Cre, 50% dilution (v2) in thalamus | Bregma AP -1.5 mm, ML ±1.0 mm, DV 3.2 mm , 100 nl | C57BL/6J | 6 light on |
| | | AAV2/9 Ef1a promoter-DIO-SN (v8) | Bregma AP 2.5 mm, ML ±1.5 mm, DV 0.8 mm , 110 nl | | |
| | Cofilin-SuperNova in ALM IT cells | AAV2/9 Ef1a promoter-DIO-CFL-SN (v7) | Bregma AP 2.5 mm, ML ±1.5 mm, DV 0.8 mm , 110 nl | TLx-Cre PL56 | 7 light on + 7 light off control |
| | Supernova- | AAV2/9 Ef1a | Bregma AP | TLx-Cre PL56 | 5 light on |

| | | | | | |
|---|---|---|---|---|---|
| | Control in ALM IT cells | promoter-DIO-SN (v8) | 2.5mm, ML ±1.5mm, DV 0.8mm , 110 nl | | |
| Fig 3 and 4 | Recording during learning | N/A | N/A | C57BL/6J | 6 light on + 2 light off |
| | Recording in expert mice | N/A | N/A | C57BL/6J or Vgat-ChR2-EYFP | 4 C57BL/6J + 6 Vgat-ChR2-EYFP, light off |
| | Recording during learning with paAIP2 manipulation in ALM $PT_{upper}$ neurons | AAVretro CamKII promoter-Cre, 50% dilution (v2) in thalamus | Bregma AP -1.5 mm, ML ±1.0 mm, DV 3.2 mm , 100 nl | C57BL/6J | 5 light on |
| | | AAV2/9 CaMKII promoter-DIO-mEGFP-P2A-paAIP2 (v3) | Bregma AP 2.5 mm, ML ±1.5 mm, DV 0.8 mm, 110 nl | | |
| | Recording during learning with paAIP2 manipulation in ALM $PT_{lower}$ neurons | AAVretro CamKII promoter-Cre, 50% dilution (v2) in Medulla | Bregma AP -6.65 mm, ML ±1.25 mm, DV 4.5 mm , 200 nl | C57BL/6J | 7 light on |
| | | AAV2/9 CaMKII promoter-DIO-mEGFP-P2A-paAIP2 (v3) | Bregma AP 2.5 mm, ML ±1.5 mm, DV 0.8 mm, 110 nl | | |
| EDF. 2 | Acute slice recording of ALM $PT_{upper}$ neurons | AAVretro CamKII promoter-Cre, 50% dilution (v2) in thalamus | Bregma AP -1.5 mm, ML ±1.0 mm, DV 3.2 mm , 100 nl | C57BL/6J (P28 at the time of injection) | 4 |
| | | AAV2/9 CaMKII promoter-DIO-mEGFP-P2A-paAIP2 (v3) | Bregma AP 2.5 mm, ML ±1.5 mm, 0.8 mm, 110 nl | | |
