## Extended Data Table 2 for "Cell-type-specific plasticity shapes neocortical dynamics for motor learning"

Extended Data Table 2. A list of viruses used in this paper

| | Virus name | Titer(vg/ml) | Packaging | ID |
|---|---|---|---|---|
| V1 | AAV2/9 CaMKII promoter-mEGFP-P2A-paAIP2 | 2.0 e13 | UNC vector core | https://www.addgene.org/91718/ CaMKIIP-mEGFP-P2A-paAIP2/pAAV was a gift from Ryohei Yasuda |
| V2 | AAVretro CamKII promoter-Cre | 3.0 e13 | Janelia viral core | https://www.addgene.org/182736/ pAAV-CamKII-Cre was a gift from Janelia Viral Tools |
| V3 | AAV2/9 CaMKII promoter-DIO-mEGFP-P2A-paAIP2 | 5.0 e12 | NA | This paper |
| V4 | AAV2/5 hsyn promoter-cre | 2.1 e13 | Addgene | https://www.addgene.org/105553/ pENN.AAV.hSyn.Cre.WPRE.hGH was a gift from James M. Wilson |
| V5 | AAV2/9 U6 promoter CaMKII gRNA-hsyn promoter-mScarlet | 1.4 e13 | NA | This paper |
| V6 | AAV2/1 hSyn promoter-mScarlet | 9.5 e12 | Addgene | https://www.addgene.org/131001/ pAAV-hSyn-mScarlet was a gift from Karl Deisseroth |
| V7 | AAV2/9 Ef1a promoter-DIO-CFL-SN | 1.1 e13 | UNC vector core | https://www.addgene.org/181740/ pAAV-EF1α-DIO-CFL-SN was a gift from Yasunori Hayashi |
| V8 | AAV2/9 Ef1a promoter-DIO-SN | 2.6 e13 | UNC vector core | https://www.addgene.org/181741/ pAAV-EF1α-DIO-SN was a gift from Yasunori Hayashi |
