## Extended Data Table 3 for "Cell-type-specific plasticity shapes neocortical dynamics for motor learning"

Extended Data Tables 3: *p-values* and sample sizes (*n)* in the paper

**Figure 1c and 2c**: *p*-Values are based on Bootstrap comparison of the mean (10000 iterations, two-sided test). α = 0.05 was divided by 4 (number of comparisons) for *Bonferroni* correction to identify significant changes (highlighted in orange in the table).

| | **Day 0** | **Day 1** | **Day 2** | **Day 3** |
|---|---|---|---|---|
| **Control** (n = 6, 7) | 0.7710 | 0.3610 | 0.3246 | 0.0424 |
| **ALM** (n = 6, 6) | 0.4956 | 0.6526 | < 0.0002 | < 0.0002 |
| **PT (both types)** (n = 5, 4) | 0.2830 | 0.2106 | 0.0048 | <0.0002 |
| **$PT_{upper}$** (n = 6, 10) | 0.6052 | 0.5516 | 0.0126 | < 0.0002 |
| **$PT_{lower}$** (n = 6, 6) | 0.8782 | 0.0294 | 0.0022 | < 0.0002 |
| **IT (layer 2/3)** (n = 6, 6) | 0.0724 | 0.3088 | 0.2888 | 0.9398 |
| **IT (layer 5)** (n = 8, 10) | 0.8240 | 0.2522 | 0.6508 | 0.8474 |

*n = (mice tested in light off condition, mice tested in light on condition)

**Figure 1f**: *p*-Values are based on Bootstrap comparison of the mean (10000 iterations, two-sided test). α = 0.05 was divided by 4 (number of comparisons) for *Bonferroni* correction to identify significant changes (highlighted in orange in the table).

| | **Day 0** | **Day 1** | **Day 2** | **Day 3** |
|---|---|---|---|---|
| N = 6, 6 | 0.8570 | 0.0440 | 0.0028 | 0.0062 |

**Figure 2e**: *p*-Values are based on Bootstrap comparison of the mean (10000 iterations, two-sided test). α = 0.05 was divided by 7 (number of comparisons) for *Bonferroni* correction to identify significant changes (highlighted in orange in the table).

| | **Day 0** | **Day 1** | **Day 2** | **Day 3** | **Day 4** | **Day 5** | **Day 6** |
|---|---|---|---|---|---|---|---|
| **$PT_{upper}$ (n = 7, 7)** | 0.2756 | 0.4916 | 0.1018 | 0.0004 | 0.0160 | 0.0078 | <0.0002 |
| **IT (layer 5) (n = 5, 5 )** | 0.5024 | 0.1770 | 0.6640 | 0.4960 | 0.0386 | 0.1004 | 0.4296 |

**Figure 2f**: *p*-Values are based on Bootstrap comparison of mean (10000 iterations, two-sided test). α = 0.05 was divided by 14 (number of comparisons) for *Bonferroni* correction to identify significant changes (highlighted in orange in the table).

| Light off | **Day 0** | **Day 1** | **Day 2** | **Day 3** | **Day 4** | **Day 5** | **Day 6** |
|---|---|---|---|---|---|---|---|

| Vs. on | | | | | | | |
|---|---|---|---|---|---|---|---|
| **$PT_{upper}$** **(n = 6, 6)** | 0.2566 | 0.3122 | 0.0004 | 0.0080 | 0.0002 | <0.0002 | <0.0002 |
| **IT (layer 5)** **(n = 5, 7)** | 0.0720 | 0.1274 | 0.1346 | 0.4684 | 0.8952 | 0.6826 | 0.4858 |

*n = (mice tested in light off condition, mice tested in light on condition)

| Cofilin vs unconjugated | **Day 0** | **Day 1** | **Day 2** | **Day 3** | **Day 4** | **Day 5** | **Day 6** |
|---|---|---|---|---|---|---|---|
| **$PT_{upper}$** **(n = 7, 6)** | 0.4672 | 0.0022 | <0.0002 | <0.0002 | <0.0002 | <0.0002 | <0.0002 |
| **IT (layer 5)** **(n = 5, 7)** | 0.7466 | 0.5442 | 0.1732 | 0.3516 | 0.9406 | 0.7716 | 0.5476 |

*n = (mice tested with Unconjugated-SN light on, mice tested with Cofilin-SN light on)

**EDF1g-i**: *p*-Values are based on Bootstrap comparison of the mean of CV or median of no response rate and # of trials (10000 iterations, two-sided test). $\alpha = 0.05$ was divided by the number of comparisons for *Bonferroni* correction to identify significant changes (highlighted in orange in the table).

| **EDF1g: CV** | **Day 0** | **Day 1** | **Day 2** | **Day 3** |
|---|---|---|---|---|
| **Control** (n = 6, 7) | 0.2720 | 0.9062 | 0.4350 | 0.8018 |
| **ALM** (n = 6, 6) | 0.1090 | 0.5972 | 0.1934 | 0.9450 |
| **PT (both types)** (n = 5, 4) | 0.0602 | 0.7384 | 0.5872 | 0.6238 |
| **$PT_{upper}$** (n = 6, 10) | 0.8830 | 0.8132 | 0.9468 | 0.5734 |
| **$PT_{lower}$** (n = 6, 6) | 0.8374 | 0.920 | 0.8216 | 0.8908 |
| **IT (layer 2/3)** (n = 6, 6) | 0.3402 | 0.3398 | 0.9950 | 0.8508 |
| **IT (layer 5)** (n = 8, 10) | 0.1508 | 0.8844 | 0.9240 | 0.2086 |

| **EDF1h: no response** | **Day 0** | **Day 1** | **Day 2** | **Day 3** |
|---|---|---|---|---|
| **Control** (n = 6, 7) | 0.4138 | 0.1396 | 0.9956 | 0.0004 |
| **ALM** (n = 6, 6) | 0.6572 | 0.0596 | 0.5542 | 0.0002 (control is higher) |
| **PT (both types)** (n = 5, 4) | 0.6572 | 0.9202 | 0.8304 | <0.0002 (control is higher) |
| **$PT_{upper}$** (n = 6, 10) | 0.4466 | 0.4642 | 0.8148 | 0.0794 |
| **$PT_{lower}$** (n = 6, 6) | 0.8322 | 0.3590 | 0.1028 | 0.0004 (control is higher) |

| | | | | |
|---|---|---|---|---|
| **IT (layer 2/3)** (n = 6, 6) | 0.8360 | 0.8132 | 0.3160 | 0.4714 |
| **IT (layer 5)** (n = 8, 10) | 0.4186 | 0.7324 | 0.4206 | 0.3476 |

| **EDF1i: # of trials** | **Day 1** | **Day 2** | **Day 3** |
|---|---|---|---|
| **Control** (n = 6, 7) | 0.2910 | 0.0162 | 0.0044 |
| **ALM** (n = 6, 6) | 0.9234 | 0.2462 | 0.3582 |
| **PT (both types)** (n = 5, 4) | 0.6254 | 0.5158 | 0.1312 |
| **$PT_{upper}$** (n = 6, 10) | 0.6482 | <0.0002 (control is lower) | 0.0178 |
| **$PT_{lower}$** (n = 6, 6) | 0.2678 | 0.0028 (control is lower) | 0.0428 |
| **IT (layer 2/3)** (n = 6, 6) | 0.0096 | 0.9254 | 0.6798 |
| **IT (layer 5)** (n = 8, 10) | 0.5308 | 0.0042 | 0.4760 |

**EDF7a-d**: *p*-Values are based on Bootstrap comparison of mean (first lick time, CV) or median no response and # of trials) (10000 iterations, two-sided test). $\alpha = 0.05$ was divided by the number of comparisons for *Bonferroni* correction to identify significant changes (highlighted in orange in the table).

| EDF7a: first lick | **Day 0** | **Day 1** | **Day 2** | **Day 3** |
|---|---|---|---|---|
| **Control vs. $PT_{upper}$** (n= 10, 5) | 0.8040 | 0.8172 | 0.0008 | <0.0002 |
| **Control vs. $PT_{lower}$** (n= 10, 7) | 0.9294 | 0.0248 | <0.0002 | 0.0002 |

| EDF7b: no response | **Day 0** | **Day 1** | **Day 2** | **Day 3** |
|---|---|---|---|---|
| **Control vs. $PT_{upper}$** (n= 10, 5) | <0.0002 (control has higher no response rate) | 0.6384 | 0.0026 (control has higher no response rate) | 0.0040 (control has higher no response rate) |
| **Control vs. $PT_{lower}$** (n= 10, 7) | 0.6762 | 0.7580 | 0.2340 | 0.0218 |

| EDF7c: CV | **Day 0** | **Day 1** | **Day 2** | **Day 3** |
|---|---|---|---|---|
| **Control vs. $PT_{upper}$** (n= 10, 5) | 0.5490 | 0.2708 | 0.4740 | 0.0496 |
| **Control vs. $PT_{lower}$** (n= 10, 7) | 0.0952 | 0.7372 | 0.1084 | 0.1750 |

| EDF7d: # of trials | **Day 0** | **Day 1** | **Day 2** | **Day 3** |
|---|---|---|---|---|
| **Control vs. $PT_{upper}$** (n= 10, 5) | 0.5128 | 0.1552 | 0.3432 | 0.1544 |
| **Control vs. $PT_{lower}$** (n= 10, 7) | 0.0260 | 0.5864 | 0.3518 | 0.8964 |

**Fig.3 and 4** Number of recorded pyramidal neurons. Only preparatory cells were analyzed.

| | | Day 1 | Day 2 | Day 3 | Expert | Sum |
|---|---|---|---|---|---|---|
| Control | # of all pyramidal cells | 244 | 270 | 265 | 1064 | 1843 |
| | # of preparatory cells | 107 | 143 | 137 | 633 | 1020 |
| $PT_{upper}$ paAIP2 manipulation | # of all pyramidal cells | 159 | 194 | 221 | 0 | 574 |
| | # of preparatory cells | 77 | 92 | 98 | 0 | 267 |
| $PT_{upper}$ paAIP2 manipulation | # of all pyramidal cells | 285 | 226 | 275 | 0 | 786 |
| | # of preparatory cells | 139 | 122 | 115 | 0 | 376 |

\# of all cells (across experimental conditions): 3203
\# of task-modulated cells (across experimental conditions): 1613

**Fig.4b:** *p*-values comparing the increase in lick time (hierarchical bootstrap; separating two PT types in a different group, unlike in the main text). $\alpha = 0.05$ is divided by the number of comparisons (in this case, 8) for *Bonferroni* correction. Since their trends are consistent, we have pooled $PT_{upper}$ and $PT_{lower}$ cells for the analysis shown in the main figure.

| | **Day 1 (within session)** | **Day 2 (within session)** | **Day 3 (within session)** | **Across 3 sessions** |
|---|---|---|---|---|
| **Control vs. $PT_{upper}$** | 0.711 | 0.002 | <0.001 | <0.001 |
| **Control vs. $PT_{lower}$** | 0.294 | 0.013 | 0.014 | 0.003 |

**Fig.4c:** p-values comparing the increase in value within sessions (hierarchical bootstrap; separating two PT types in a different group, unlike in the main text):

| | **Day 1 (within session)** | **Day 2 (within session)** | **Day 3 (within session)** | **Across 3 sessions** |
|---|---|---|---|---|
| **Control vs. $PT_{upper}$** | 0.648 | 0.144 | 0.008 | <0.001 |
| **Control vs. $PT_{lower}$** | 0.301 | 0.095 | 0.023 | 0.004 |

**Fig.4d:** p-values comparing the decrease in value within sessions (hierarchical bootstrap; separating two PT types in a different group, unlike in the main text).

| | **Day 1** | **Day 2** | **Day 3** |
|---|---|---|---|
| **Control vs. $PT_{upper}$** | 0.25 | 0.003 | 0.273 |

| | | | |
|---|---|---|---|
| **Control vs. $PT_{lower}$** | 0.349 | 0.021 | 0.512 |
