## Extended Data Table 4 for "Cell-type-specific plasticity shapes neocortical dynamics for motor learning"

Extended Data Table 4. A list of parameters used for the network model
See Methods for descriptions of parameters.

| Parameter name | Value (synaptic potentiation model) | Value ( synaptic depression model) |
|---|---|---|
| $\tau_{PT} = \tau_{IT}$ | *10 ms* | *10 ms* |
| $\tau_{I}$ | *10 ms* | *10 ms* |
| $\tau_{Th}$ | *10 ms* | *10 ms* |
| $W_{IT \leftarrow IT}$ | *1.22* | *1.02* |
| $W_{PT \leftarrow IT}$ | *0.41* | *0.5* |
| $W_{In \leftarrow IT}$ | *0.5* | *0.55* |
| $W_{PT \leftarrow PT}$ | *0.6* | *0.8* |
| $W_{In \leftarrow PT}$ | *0.6* | *0.65* |
| $W_{Th \leftarrow PT}$ | *0.6* | *0.55* |
| $W_{In \leftarrow In}$ | *1.45* | *0.43* |
| $W_{IT \leftarrow In}$ | *0.6* | *0.55* |
| $W_{PT \leftarrow In}$ | *0.6* | *0.65* |
| $W_{IT \leftarrow Th}$ | *0.3* | *0.8* |
| $W_{PT \leftarrow Th}$ | *0.6* | *0.8* |
| $W_{In \leftarrow Th}$ | *1.1* | *0.4* |
| $b_{IT}$ | *0.35* | *0.1* |
| $b_{PT}^{max}$ | *1.2* | *NA* |
| $b_{PT}^{min}$ | *NA* | *0.26* |
| $b_{In}$ | *0.2* | *0.4* |
| $b_{Th}$ | *0* | *0* |
| $c_{IT}$ | *0* | *0* |
| $c_{PT}$ | *0* | *0* |
| $c_{In}$ | *0* | *0* |
| $c_{Th}$ | *0.75* | *0.75* |
| $I_{IT}$ | *-0.1* | *-0.1* |

| | | |
|---|---|---|
| $\boldsymbol{I_{PT}}$ | *-0.1* | *-0.1* |
| $\boldsymbol{I_{In}}$ | *-1.5* | *-1.0* |
| $\boldsymbol{I_{Th}}$ | *-0.5* | *-0.2* |
| $\boldsymbol{I^{tonic}}$ | $0.85 * \frac{\log(k+1)}{\log(7)}$, for k=0,…,6 | $-0.11 * \frac{\log(k+1)}{\log(7)}$, for k=0,…,6 |
| $\boldsymbol{r^{*}_{PT}}$ | *15* | *15* |
| $\boldsymbol{b^{0}_{PT}}$ | *0* | *0.7* |
| $\boldsymbol{\mu^{tonic}}$ | *0* | *0* |
| $\boldsymbol{\sigma^{tonic}}$ | *0.2* | *0.05* |
| $\boldsymbol{\mu^{cue}}$ | *3.0* | *5.0* |
| $\boldsymbol{\sigma^{cue}}$ | *0.35* | *0.25* |
| $\boldsymbol{I^{cue}}$ **duration** | *100 ms* | *100 ms* |
| $\boldsymbol{\eta}$ | *0.00075* | *0.00025* |
